# CODAvision: best practices and a user-friendly interface for rapid, customizable segmentation of medical images

**DOI:** 10.1101/2025.04.11.648464

**Authors:** Valentina Matos-Romero, Jaime Gómez-Becerril, André Forjaz, Lucie Dequiedt, Tyler Newton, Saurabh Joshi, Yu Shen, Eban Hanna, Praful Nair, Arrun Sivasubramanian, Jenny S. H. Wang, Emily L. Lasse-Opsahl, Alexander T.F. Bell, Julianna Czum, Charles Steenbergen, Dao-Fu Dai, Laura D. Wood, Luciane T. Kagohara, Elana J. Fertig, Marina Pasca di Magliano, Joseph J. Shatzel, Owen J.T. McCarty, Jamie O. Lo, Avi Rosenberg, Ralph H. Hruban, Arrate Muñoz-Barrutia, Denis Wirtz, Ashley L. Kiemen

## Abstract

Image-based machine learning tools have emerged as powerful resources for analyzing medical images, with deep learning-based semantic segmentation commonly utilized to enable spatial quantification of structures in images. However, customization and training of segmentation algorithms requires advanced programming skills and intricate workflows, limiting their accessibility to many investigators. Here, we present a protocol and software for automatic segmentation of medical images guided by a graphical user interface (GUI) using the CODAvision algorithm. This workflow simplifies the process of semantic segmentation of microanatomical structures by enabling users to train highly customizable deep learning models without extensive coding expertise. The protocol outlines best practices for creating robust training datasets, configuring model parameters, and optimizing performance across diverse biomedical image modalities.

CODAvision enhances the usability of the CODA algorithm (*Nature Methods*, 2022) by streamlining parameter configuration, model training, and performance evaluation, automatically generating quantitative results and comprehensive reports. We expand beyond the original implementation of CODA to serial histology by demonstrating robust performance across numerous medical image modalities and diverse biological questions. We provide sample results in data types including histology, magnetic resonance imaging (MRI), and computed tomography (CT). We demonstrate the diverse use of this tool in applications including quantification of metastatic burden in *in vivo* models and deconvolution of spot-based spatial transcriptomics datasets. This protocol is designed for researchers with interest in rapid design of highly customizable semantic segmentation algorithms and a basic understanding of programming and anatomy.

## Introduction

Deep learning has emerged as a powerful tool for analyzing digitized biomedical images in various research applications.^1–11^ Semantic segmentation models are commonly used due to their ability to be quickly adapted to specific datasets for detecting structures across a variety of image types.^12–14^ In addition, their relatively smaller architectures (compared to some larger transformer and foundation models) make them amenable to retraining by researchers without access to high-performance computing. However, the implementation of these methods often requires advanced programming skills and intricate workflows, limiting their accessibility and widespread adoption outside of the computational biology community. In response, some popular deep learning workflows have been made more accessible through the development of graphical user interfaces (GUIs), which reduce or eliminate the need for extensive coding during implementation.^15–19^

To address this challenge, we developed a novel GUI aimed at simplifying the process of training highly-customizable models for segmentation of biological structures in medical images. As a companion to this interface, we developed an extensive guide of best practices for annotation layer selection and annotation style for the construction of robust supervised models.

### How this Protocol Improves Upon the Existing CODA Methodology

This protocol is a powerful extension of the CODA workflow.^20^ CODA, a MATLAB-based pipeline for reconstruction of serial histological images into quantitative 3D datasets, has been used extensively in biomedical research applications including study of pancreatic cancer progression, heart development, diabetic neuropathy, and skin regeneration, among others.^20–42^ CODA has shown technological power beyond its original presentation as a method to create 3D maps from serial hematoxylin and eosin (H&E) stained histology. Numerous recent studies have utilized one useful module of CODA, its image segmentation workflow, and integrated it with spatially resolved genomics,^21^ spatial transcriptomics and proteomics,^30–32^ tissue stiffness,^33^ antibody-based staining techniques,^34–36^ organoid modelling,^37,38^ and to quantify *in vivo* histology.^39–41^ Here, we extract the segmentation module of the CODA package and dramatically improve its speed, usability, and applicability to diverse biomedical image modalities. In the studies described earlier the core implementation codes of CODA remained unchanged. This original implementation possess three clear limitations which we address in the current protocol:

(1) The original package is written as discrete functions in MATLAB. This language is not open source, and the format of the codebase made its implementation challenging even for users with extensive programming experience. We address this through the translation of the CODA segmentation workflow to Python, optimization of the code speed and performance, and creation of a user-friendly GUI. We name this optimized workflow CODAvision. (2) The original package lacked description of the format and style of manual annotations required for training robust segmentation models. Recent groups have highlighted the importance of training dataset quality in deep learning approaches.^43,44^ We address this through generation of extensive user guides describing best-practices for rapid generation of robust segmentation models. (3) The original package demonstrated applicability to H&E images only. Here, we demonstrate applicability of CODAvision to H&E as well as other imaging types including MRI and CT.

The novelty of this workflow is its depth in describing best practices for the optimal creation of training datasets, construction of an intuitive user interface for parameter configuration and model architecture selection, and automatic generation of model performance reports and quantitative results for streamlined use in scientific experiments (**Fig 1A**). For example, using a dataset of mouse lung histology we demonstrate the ability of CODAvision to rapidly generate quantitative spatial measurements of composition from *in vivo* experiments. To validate CODAvision’s capability to quantify metastatic burden in in vivo studies, we analyzed 55 H&E-stained lung tissue sections from mice injected with MDA-MB-231 breast cancer cells. Using a DeepLabv3+ architecture with a ResNet50 backbone (precision and recall >90% for all tissue types), we quantified the metastatic coverage, which revealed 45% and 53% metastatic burden for wild-type and scrambled control cells, respectively (**Fig 1B**).

**Figure 1:**
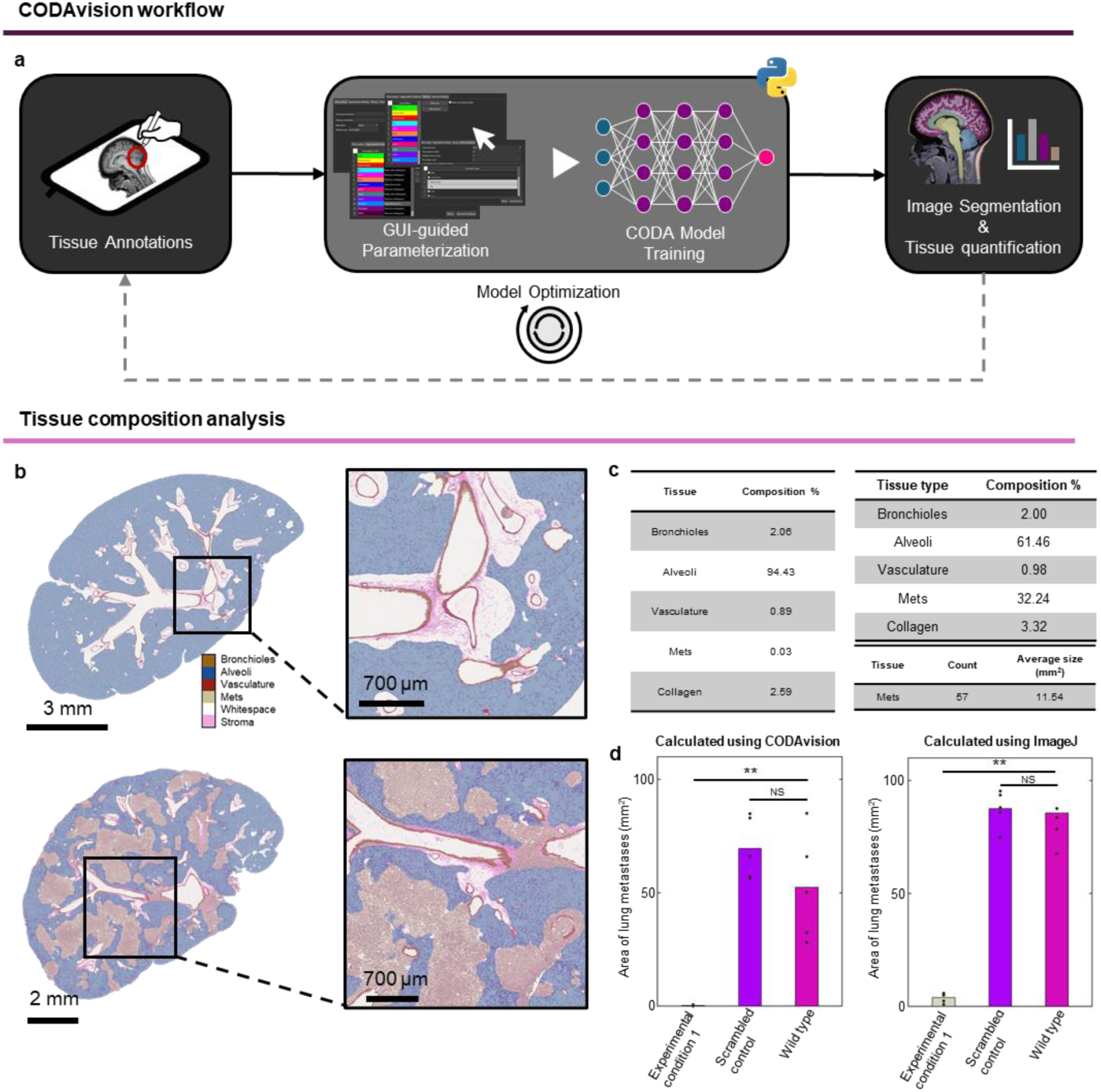
CODAvision workflow and sample application. a. Pipeline overview showing sequential steps: tissue annotation for dataset creation, GUI-guided parameterization, model training, and quantitative analysis. b. Representative semantic segmentation results comparing lung histology from a mouse used in a control arm (top) to histology from a mouse used in the experimental arm (bottom) of an *in vivo* experiment. c. Sample tissue composition analysis with metastases object count for sample b (bottom). d. Comparison of metastatic burden occupied in the lung across three experimental conditions, showing CODAvision quantification (left) versus a conventional ImageJ-based analysis (right).

### Comparison to Alternate Techniques

Compared to alternative techniques for quantifying structures in biomedical images, our workflow possesses differences that make it advantageous for certain research questions. In the proposed protocol, we guide users through the generation of robust datasets to train customizable deep learning models. Another approach is use of large-scale foundation models and vision transformers that have been trained on millions of examples for object-detection tasks, or sometimes more specifically on extraction of meaningful features from biomedical images.^8,45–48^ These methods differ from our protocol in that they are intended to be fully automated, where our workflow requires initial manual annotation.

For research projects where direct measure of a certain structure (such as cancer metastases in mouse histology, or something very specific such as a subtle phenotype of cancer cells that is not currently well defined) is desired, our workflow enables rapid segmentation of that specific anatomical structure. In contrast, for applications that benefit from a more holistic perspective – such as generalized feature extraction or the generation of attention maps for survival prediction – pre-trained foundation models may be more suitable. Our workflow is designed to run on a standard desktop computer and requires only minimal programming expertise, making it broadly accessible. On the other hand, the deployment of large foundation or transformer models typically demands high-performance computing resources and advanced computational skills.

### Experimental design

In this section, we provide a methodological overview of the five main steps outlined in the protocol: dataset creation, GUI-guided parametrization, model training, model optimization, and image classification using a custom pretrained model.

Example datasets: To help users explore the CODAvision software and its features, we include a link to a sample dataset of annotated mouse lung histology. Results obtained from CODAvision analysis of these datasets are presented in the Anticipated Results below. We recommend that users initially run CODAvision on the supplied dataset and follow the detailed protocol described in the Procedure section before analyzing new data.

### Dataset creation

A dataset for this workflow consists of a set of biomedical images (e.g., digitized histology, MRI, CT.) and their associated annotation metadata. The first step of the workflow is to identify structures in the dataset that can be distinguished from each other, and which of those are required for the research objective. An exhaustive list of structures is then made, and manual these structures annotated in the freely available program Aperio ImageScope. The selection of classes directly corresponds to the research question the CODAvision analysis aims to address (BOX 1). In Procedure 1, we outline strategies for developing a robust training dataset that enables building custom segmentation models for the targeted structures.

### GUI guided parametrization

Once the manual annotations are complete, the next step in the CODAvision workflow is to define the date import and model training parameters using a Python-based GUI. Users first install the CODAvision Python package, following the instructions for codes and dependencies outlined in Procedure 2. The GUI guides users through configuring settings for model training, including specifying the location of training and testing datasets, selecting an image resolution (downsampled files are generated using OpenSlide),^49^ and customizing model parameters. The choice of resolution is critical, balancing segmentation detail with computational efficiency. The GUI further allows for the management of the manual annotated layers, enabling features such automatic removal of background pixels, combining or deleting annotation layers, and defining the nesting logic for managing overlapping annotations. Advanced settings enable further customization, such as adjusting tile size, batch size, and model architecture (e.g., DeepLabV3+, UNet),^12,50^ allowing users to optimize training based on their computational resources and dataset size. Once all parameters are set, the model will begin training.

### Model training

The model training phase begins with tissue thresholding, guided by an interactive popup window that allows users to fine-tune the threshold cutoff for optimal tissue/background separation. Once the threshold is set, the pre-processing and model training proceed automatically. During this phase, the .xml annotation coordinate data will be imported and converted to .png annotation masks. These masks will be used to create training and validation tiles built using data augmentation techniques such as hue adjustment, scaling, rotation, and Gaussian filtering to enhance dataset heterogeneity and model robustness. The chosen model architecture is trained on these augmented tiles. After training, the model performs inference on test images, generating a confusion matrix that showcases the precision, recall, and overall accuracy. For each segmented image, the workflow will output a classified .png mask and a colorized .jpg overlay, enabling users to rapidly review the model performance using both quantitative (confusion matrix) and qualitative (overlay images) review. The overall composition of each image will be saved in a .csv file, along with more detailed morphological calculations if desired by the user. The model results will be automatically summarized in a generated .pdf report. Users are encouraged to review these results to ensure the trained model meets the recommended performance benchmarks (e.g., >90% overall accuracy and >85% per-class precision and recall, and visually acceptable results).

### Model optimization

If the performance of the trained model is unsatisfactory, Procedure 4 outlines steps to efficiently retrain and optimize the model. Optimization strategies include review of the colorized mask overlay images to identify patterns of misclassification, determining if any structures were missed during initial annotation, and adjusting model parameters to better align with the desired results. If misclassifications persist, it is recommended to expanded the training dataset by adding new images rather than adding annotations to existing training images. These optimization steps can be repeated iteratively until satisfactory results are achieved, ensuring the model meets the recommended performance benchmarks.

### Image segmentation with custom pretrained model

Once a model is trained, users can apply it to segment additional images beyond the original training set using a pretrained model. The steps for classifying additional images through \the CODAvision GUI are detailed in the Procedure 5. Users can select the images to segment, choose a pretrained model for inference, and optionally modify the color palette for the colorized segmentation masks or perform additional object-based morphology analysis.

### Description of the Expertise Needed to Implement the Protocol

This protocol is designed for researchers with some training and experience in computational biology. In particular, users must possess some knowledge of anatomy and of coding to use this workflow, which we describe here.

Successful implementation of this workflow requires knowledge of anatomical structures and how they appear in histology / radiology images. For example, a user wishing to segment cancer metastases in H&E images of mouse lung must understand how to differentiate cancer cells from the functional cells of the lung in these images. The user will use this knowledge to manually annotate and to qualitatively assess model performance. We provide a supplemental annotation guide (**supplementary file 1**) that gives some background on how to identify structures in medical images, which may be beneficial to some users.

Users must also possess a basic understanding of programming. While operation of this workflow is streamlined with a user-friendly GUI, initial installation requires familiarity with Python scripting, CUDA, and cuDNN setup for GPU acceleration. Detailed instructions for these steps are provided on the GitHub page, where all codebase and dependencies are hosted. We suggest that users without programming knowledge obtain assistance when initially installing the package, after which operation of the GUI can proceed without significant coding expertise.

### Limitations

This workflow, while powerful, possesses several limitations which we document here. First, some the model architectures included (DeeplabV3+ and U-Net) may require significant computational resources, such as high-performance GPUs. The speed of this workflow is significantly impacted on computers without GPUs or on standard laptops.

Second, the segmentation models described here require highly specific manual annotations for training data. The benefit of this approach is the ability to, in the span of a few days, train highly accurate and highly customizable segmentation models in any cohort. The limitation is that a model trained on one organ (for example mouse lungs) is not easily adapted to another organ (for example human pancreas). Instead, users must generate new manual annotations for each new application.

## Materials

### Equipment

1. computer with at least 16 GB of RAM
2. NVIDIA graphics processing unit (GPU) with at least 8 GB RAM
3. up-to-date operating system (Windows 10/11, OSX 11)
4. least 2.5 GB of storage space
5. working CUDA (≥11.2) and CuDNN (≥8.1) installation (instructions available at the provided GitHub page) In the analysis described here we used a computer with the following specifications:
6. with 128 GB RAM and an NVIDIA GeForce RTX 4090 GPU running on Windows 11.

### Software

1. CODAvision software available in the following repository: https://github.com/Kiemen-Lab/CODAvision.
2. Python Interpreter (e.g., PyCharm, Visual Studio, Spyder)
3. Image Annotation Tool: For annotating images, the Aperio ImageScope application is required. This software can be installed from the following link: https://www.leicabiosystems.com/digital-pathology/manage/aperio-imagescope.
4. Example dataset:

**Dataset 1**: Mouse lung histology, available at the following link: https://drive.google.com/drive/folders/1K-wY_ArVGbEhebQD4AjOeERwx6-4Fw3G.

## Procedure

CRITICAL: This protocol assumes that users possess a dataset of images intended for semantic segmentation and quantification. For sample datasets, see Datasets 1 and 2 in the software section of this protocol. We demonstrate that CODAvision can be applied to diverse image types including histology, MRI, and CT. Our primary demonstration, and the sample datasets provided are histological images scanned at 20× magnification (roughly 0.5 µm / pixel resolution), though the procedure could be similarly applied to images scanned at higher or lower resolution.

### Procedure 1: Constructing a training dataset for deep learning training

#### Annotating on Aperio ImageScope

##### • Timing: 10 - 20 h

1. Select six images to annotate from the initial cohort. CRITICAL: The images selected for annotation should reflect the heterogeneity of the larger dataset. This may include selecting images from different scientific groups (control vs experimental conditions), images possessing distinct anatomical features, and images with technical heterogeneity such as variation in lighting or focus. Construction of a heterogeneous training dataset will improve the robustness of the segmentation.
2. Create a folder named ‘Training dataset’, and another folder named ‘Testing dataset.’
3. Copy five of the selected images to the ‘Training dataset’ folder and save the sixth image in the ‘Testing dataset’ folder.
4. Install Aperio ImageScope following the installation instructions available at: https://www.leicabiosystems.com/digital-pathology/manage/aperio-imagescope.
5. We suggest users change two settings in Aperio Imagescope upon installation to improve the user experience.
  a. To increase the maximum allowable zoom for precise annotation, navigate to Tools > Options > General Tab > Maximum magnification and enter 1000%.
  b. To automatically save annotations when exiting the program, navigate to Tools > Options > Annotation Tab > Annotation Settings, and check the box ‘Automatically save annotation changes’.
6. Open one of the images from the ‘Training dataset’ folder in Aperio ImageScope.
7. Create the annotation layers by navigating to View, then Annotations to show the ‘Annotations - Detailed View’ window and click the ‘+’ button to add an annotation layer. Rename the annotation layer by clicking on the layer name’s top and press ‘F2,’ then input the desired name.
8. Create one annotation layer for each object you would like to train a model to segment. Once all layers are created, press save. This will generate an .xml file corresponding to this image that will contain the annotation coordinates that will be used for model training. CRITICAL: All annotated images must have the same layer order. Create a layer for every structure, even in images where those structures are absent. CRITICAL: The testing dataset must contain at least one annotation of each annotation layer. If all layers are not present in any single image, consider using multiple images for testing so that the overall testing dataset contains at least one annotation of each annotation layer. CRITICAL: The .xml files in the training and testing folders must correspond to the images they annotate, with identical filenames. If an .xml file has a different name than its associated image, the code will fail. Ensure that every image has a matching .xml file and vice versa. **How to choose which structures to annotate? BOX 1** The number of annotation layers you should generate depends on your research objective. In histology, many cell types can be differentiated by the trained eye, including various epithelial, vascular, and stromal compartments. We suggest that users first make a list of the major structures present in their images, then group these structures until the desired granularity is obtained. See **Fig 2** for an example of a high-detail and a low-detail model trained on fetal rhesus macaque kidney histology. Where exhaustive anatomical labelling is desired, the user can generate a highly specific list of structures identifiable in H&E (**Fig 2A**). For a more focused project, the user can group labels to reduce the number of annotation layers and increase the speed of the project (**Fig 2B**). CRITICAL: No matter your research question, all models should contain a background or whitespace layer in the annotation dataset to contain non-tissue pixels in the image.
9. Begin annotating using the ‘Pen’ tool. Start the annotation by clicking and annotating the area of interest in the main window. Use the pen tool to manually outline the region of interest, closing the drawn shape once complete. If you wish to modify the annotation, click and redraw over the annotated region until you achieve the desired level of precision in refining the borders. CRITICAL: The resulting quality of your segmentation model relies on the quality of your annotations. Zoom in to high magnification when annotating and aim to annotate the structure boundaries very cleanly and consistently (**Fig 3B**). CRITICAL: We recommend that users periodically save the annotations manually by clicking the save button in the ‘Annotations - Detailed View’ window. This will help prevent potential data loss.
10. Make ∼20 annotations per tissue structure per image (training and testing). For rare structures, there may be fewer than 20 examples per image. <u>GOOD annotation</u> practices: here we provide guidance on good annotation practices. For more detailed notes on annotating, see the companion Annotation Guide in **supplementary file 1**.
11. Nesting: make overlapping annotations of different classes following a consistent nesting hierarchy (**Fig 3A**). CRITICAL: Define the nesting hierarchy before beginning annotations. This hierarchy must remain consistent across all annotated images and will be used during the deep learning model parametrization described in Procedure 2. **What is nesting? BOX 2** We have observed that CODA segmentation models yield better results when the tissues are identified within their microenvironments. To achieve this, employ an annotation technique called ‘nesting.’ Nesting uses a hierarchical tissue organization, in which higher level tissues can be ‘nested’ inside lower-level ones. **Fig 3A** illustrates the arrangement of three hypothetical annotated types: tissue A (triangles), tissue B (squares), and tissue C (circles). Varying the nesting hierarchy changes how these annotation layers are imported for model training.
12. Annotate structures across the entire image, not just in one region.
13. Include diverse examples of all annotation classes. Building a dataset with varied morphologies for each class ensures optimal performance of your model on unseen data.
14. Include ‘non-ideal’ annotations for each class to enable the model to correctly classify tissue types even in the presence of noise. For example, annotate structures that are slightly blurry, darker, or paler.
15. When glandular structures that contain a lumen, include background annotations within the lumen if noise, such as fluid or red blood cells, is present (**Fig 3A**, right).
16. Include 5-10 annotations at tissue structure edges when annotating whole slide images to ensure accurate differentiation between tissue borders and background during classification.
17. When re-annotating to improve model performance, first review the classified images to identify regions of misclassification. Focus your annotations on these regions to efficiently correct the model (**Fig 3C**). CRITICAL: Refer to the supplemental annotation guide (**supplementary file 1**) for detailed examples of tissue annotations and best practice.

### Procedure 2: Defining model parameters using the GUI

#### Timing: 5 min

1. After completing the annotation step, download CODAvision by following the README instructions present on the following GitHub page: https://github.com/Kiemen-Lab/CODAvision.
2. If desired, import the Python code to an IDE (Integrated Development Environment).
3. Run the CODAvision.py code to execute the GUI to parametrize the settings for the model. <u>File Location tab (</u>**Fig 4A**)
4. Browse for the folder containing the training annotations. This folder should contain the annotated images and .xml files generated during Procedure 1.
5. Repeat step 4 to browse for the folder containing the testing annotations.
6. Specify a desired image resolution by selecting an option from the dropdown list. **How to choose a training resolution? BOX 3** The choice of training resolution is critical for achieving the desired segmentation accuracy while managing computational resources efficiently. For cellular-level analyses, we recommend using 10× (1 μm/pixel), whereas organ-level or large tissue structures can be effectively segmented at 1× (8 μm/pixel). The key consideration is the trade-off between segmentation detail and computational efficiency. Higher resolutions provide finer detail but result in larger file sizes, extended processing time, and require higher precision manual annotations. CODAvision also enables users to provide pre-generated downsampled images and to input this custom scale factor instead of choosing from one of the predefined resolutions. To do this, select “custom scale” and browse for the folder containing the scaled .tif or .png images.
7. Enter a desired name for the deep learning model. By default, the name is prepopulated with today’s date, but may be customized.
8. (Optional) Select ‘Custom’ from the ‘Resolution’ dropdown menu to train on resolutions different from the default options. This action will display a scaling factor input field where you can specify the desired downsampling ratio (must be ≥1) (**Fig 5**).
  a. To use different images for downsampling instead of the annotated images, select the ‘Scale custom images’ checkbox. Then, locate the custom image directories, which should be organized into separate training and testing folders. The image filenames must correspond exactly to their respective annotation files.
  b. If your custom images are pre-scaled, activate the ‘Use pre-scaled images’ checkbox. Ensure that these images match the value specified in the ‘Scaling factor’ field and are .tif or .png filetype.
  CRITICAL: Choose this option when working with images not originally annotated in the recommended formats (.ndpi or .svs). The custom image downsampling feature accepts the following file formats: .ndpi, .svs, .tif, .jpg, .png, and .dcm.
9. After completing all sections on the ‘File Location’ tab, click ‘Save & Continue’ move to tab 2: ‘Segmentation Settings.’ This tab serves as the interface for defining the different classes for the deep learning model, as well as inputting nesting information. This page will be pre-populated with the annotation classes and colors used during the manual annotation step in Aperio ImageScope. <u>Segmentation Settings tab (</u>**<u>Fig 4B</u>**)
10. Determine how to handle the white/background pixels (referred to as ‘whitespace’ in this protocol) for each annotation layer. To do so, click on an annotation class from the table, and select one of the three available options in the ‘Annotation class whitespace settings’ (see **Fig 6** for examples) **How to manage background in manual annotations? BOX 4** Proper whitespace management is critical for training high-accuracy deep learning models. Here, we provide advice on what when to automatically remove whitespace or non-whitespace from your annotation layers: Select ‘Remove whitespace’ to eliminate the background pixels from an annotation layer. This is relevant for scenarios such as excluding the lumen from a glandular structure or to remove the white pixels intermixed between stromal fibers. Select ‘Keep only whitespace’ to retain only the background pixels in the annotation. This is relevant when annotating fat and aiming to exclude nonwhite lines separating individual fat cells. Select ‘Keep tissue and whitespace’ to retain both background and nonwhite pixels. This is appropriate for the noise/background layer, as the annotated regions may contain both whitespace and noise such as shadows, or when annotating a solid structure such as hepatocytes in the liver. Refer to **Fig 6** for visual examples.
11. Upon selecting an ‘Annotation class whitespace settings’ option, click ‘Apply’ or ‘Apply all’ to update the table.
12. Define the destination class for whitespace pixels removed from annotation layers where the option ‘Remove whitespace’ was selected. In general, the destination class should be the background class.
13. Similarly, define the destination class for removed non-whitespace pixels taken from the annotation layers where the option ‘Keep only whitespace’ was selected. In general, the destination class should be the stromal class. This input must be defined even if no annotation layer was assigned with ‘Keep only whitespace.’
14. (Optional) To change the color assigned to any annotation layer, select the desired annotation class from the table click the ‘Change Color’ button. In the color picker window, select the desired color, then Click ‘OK’ to confirm the color change. CRITICAL: By default, the model’s classification output will use the same colors as those used during manual annotation (shown as background colors in the table). Color changes are purely aesthetic and do not affect model performance, but well-chosen colors may improve users’ ability to visually interpret and present the segmentation results. To ensure accessibility, we recommend selecting color palettes that are color-blind friendly.
15. (Optional) To combine annotation layers, hold the ‘ctrl’ key and select the desired rows in the table. Click the ‘Combine classes’ button and, when prompted, enter a name for the combined class. In the color picker window, select a color for the combined class.
16. (Optional) To delete unwanted annotation layers, select the appropriate row by clicking inside the table and click the ‘Delete class’ button. CRITICAL: Delete unwanted annotation layers that should not be included the model training. This is useful for removing empty layers.
17. (Optional) Click the ‘Reset list’ button to return the annotation class table to its default state. This will remove whitespace management choices, uncombine any combined layers, and restore deleted layers.
18. Once this tab is completed, click ‘Save & Continue’ in the bottom right corner. The interface will automatically advance to tab 3: ‘Nesting’. <u>Nesting tab (</u>**Fig. 4C**)
19. Define the appropriate nesting order to ensure correct handling of overlapping annotation classes. This order should have been established during the manual annotation step in Procedure 1. To define the nesting order in the GUI, select a desired annotation layer. Use the ‘Move Up’ and ‘Move Down’ buttons to adjust its position according to layering priority. CRITICAL: The ‘Nesting tab’ allows the user to configure the layering hierarchy of annotation classes from lowest to highest priority in cases of overlap. The class at the top of the table are assigned the highest priority, while the class at the bottom are lowest priority.
20. (Optional) To define different nesting orders to several annotation layers that combined in the Segmentation Settings Tab, check the ‘Nest uncombined data’ box. CRITICAL: If annotation classes were combined in the previous tab, the nesting table will display these combined layers by default. If a combined layer is made from two layers that require different nesting priority, check the ‘Nest uncombined data’ box. Refer to the annotation guide in **supplementary file 1** for detailed examples on establishing the nesting order.
21. Once the nesting tab is completed, the user has four options:

a. (Most likely) Select ‘Save and train’ to immediately proceed with model training
b. Click ‘Save and close’ to save the model configuration but NOT train the model.
c. Click ‘Continue to advanced settings’ to define more complex model parameters before training the model. This tab will enable the user to adjust model hyperparameters, select the model architecture, or select classes for advanced quantitative analysis.
d. Select ‘Return’ to go back to Tab 2. <u>Advanced Settings tab (</u>**Fig. 4D**)
22. (Optional) To adjust the default tile size input to the segmentation model, choose a new tile size from the dropdown list (default: 1024 × 1024 RGB). CRITICAL: The training tile size default is 1024 × 1024 pixels. Users with GPU RAM constraints should consider a smaller size such as 512 × 512 or 256 × 256 (the size must be a power of 2).
23. (Optional) To adjust the training and validation tile number, click on the up and down arrows next to each respective text box. The default number of training tiles is 15 and the default number of validation tiles is 3. Users may increase these numbers for especially large (>50 annotated images) or small (<5 annotated images) training datasets.
24. (Optional) Adjust the number of images used for tissue mask thresholding (described in greater detail in Procedure 3) by clicking on the up and down arrows next to the ‘Tissue mask evaluation:’ text box. By default, this number is three images, but a higher number may be desired for larger or more diverse datasets.
25. (Optional) Click the ‘Individual tissue mask evaluation’ checkbox to customize the tissue mask threshold for each image in the dataset. This option is recommended when image appearance varies substantially across the dataset and images may have varying background intensity thresholds.
26. (Optional) Choose the desired model architecture from one of two available options in the dropdown box (DeepLabV3+, UNet). By default, the training architecture is DeeplabV3+. **How to choose a model architecture? BOX 5** The choice of model architecture depends on computational resources available and the specific structures to be segmented. UNet contains approximately 41 million parameters, and in our benchmark test required approximately 40 minutes to train on an NVIDIA GeForce RTX 4090 GPU. In contrast, DeepLabV3+ contains approximately 12 million parameters, and trained in approximately 75% of the training time of UNet. While UNet excels in capturing fine-grained details due to its deeper architecture, DeepLabV3+ is often more computationally efficient and may be preferable for users with limited computational resources or time constraints. Users are encouraged to experiment with both architectures to determine which best suits their specific needs. For computationally experienced users, additional architectures can be integrated into the workflow by modifying the codavision/models/backbones.py code, where the provided networks are implemented as Python classes. Users can also adapt the codavision/models/training.py and codavision/CODA.py to enable training on the new architecture and include it as an option in the dropdown menu of the advanced settings tab in the GUI.
27. (Optional) Modify the batch size used for training. By default, the batch size is set to 3. CRITICAL: Adjust the batch size according to your GPU memory capacity. Larger batch sizes accelerate training but require more GPU memory, while smaller batch sizes increase training duration but reduce memory requirements. Monitor GPU memory usage during initial training attempts to optimize this parameter for your system.
28. (Optional) Select classes for detailed quantitative analysis (object number count and size per image). CRITICAL: For a selected class in each segmented image, the total number of objects >500 pixels will be counted and the object size in pixels will be documented. These data will be exported in a .csv file following model training and image segmentation.
29. nce the advanced settings tab is completed, the user has three options:

a. (Most likely) Select ‘Save and train’ to proceed with model training.
b. Select ‘Save and close’ to save the configuration but NOT train the model.
c. Select ‘Return’ to go back to tab 3.

**Figure 2:**
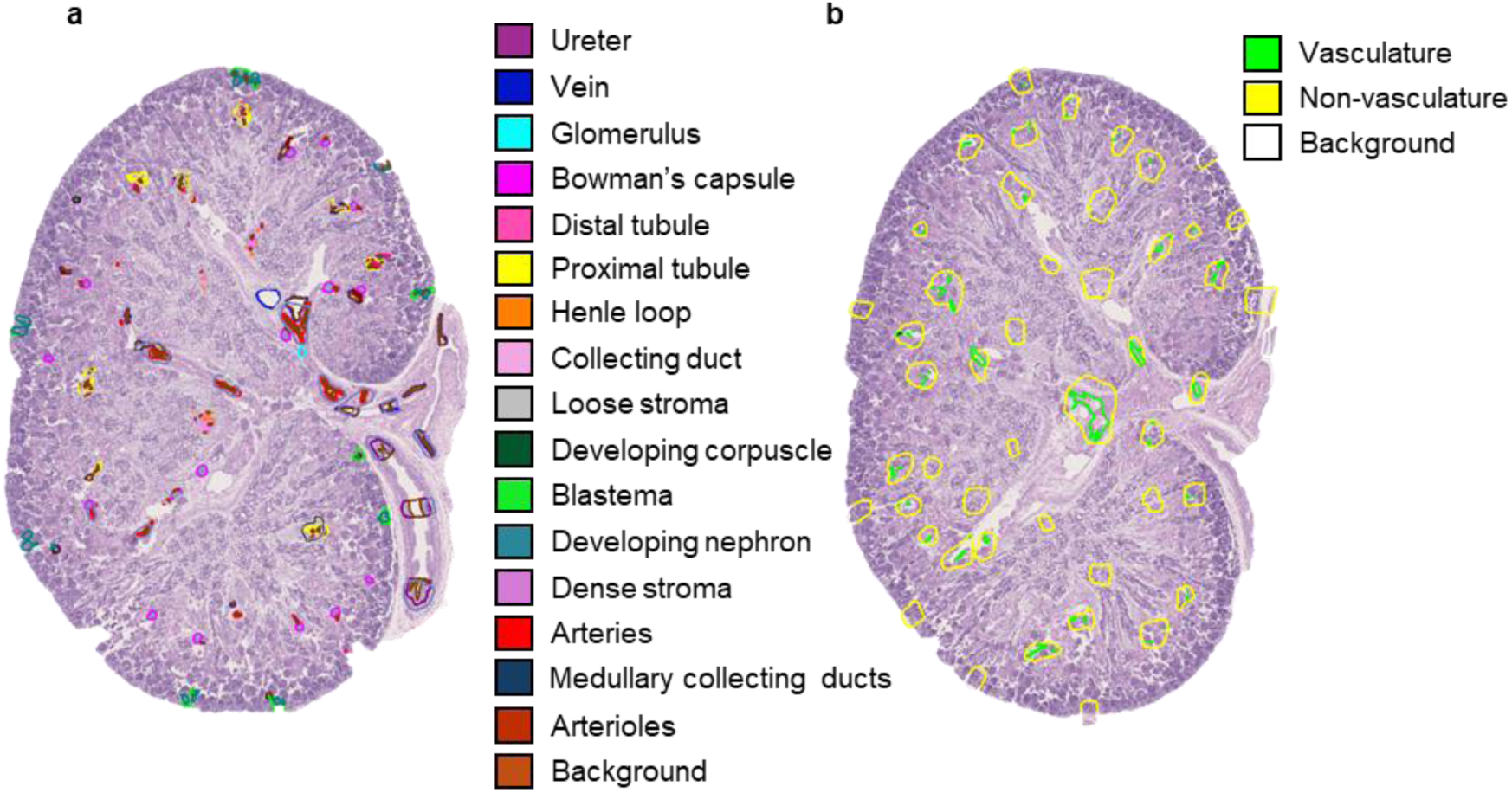
Sample histological image containing fetal rhesus macaque kidney with anatomical annotations overlaid. a. For a high-detail model, seventeen tissue structures are identifiable in the kidney. b. For a vasculature-focused model, the annotation layers can be grouped to remove unnecessary labels.

**Figure 3:**
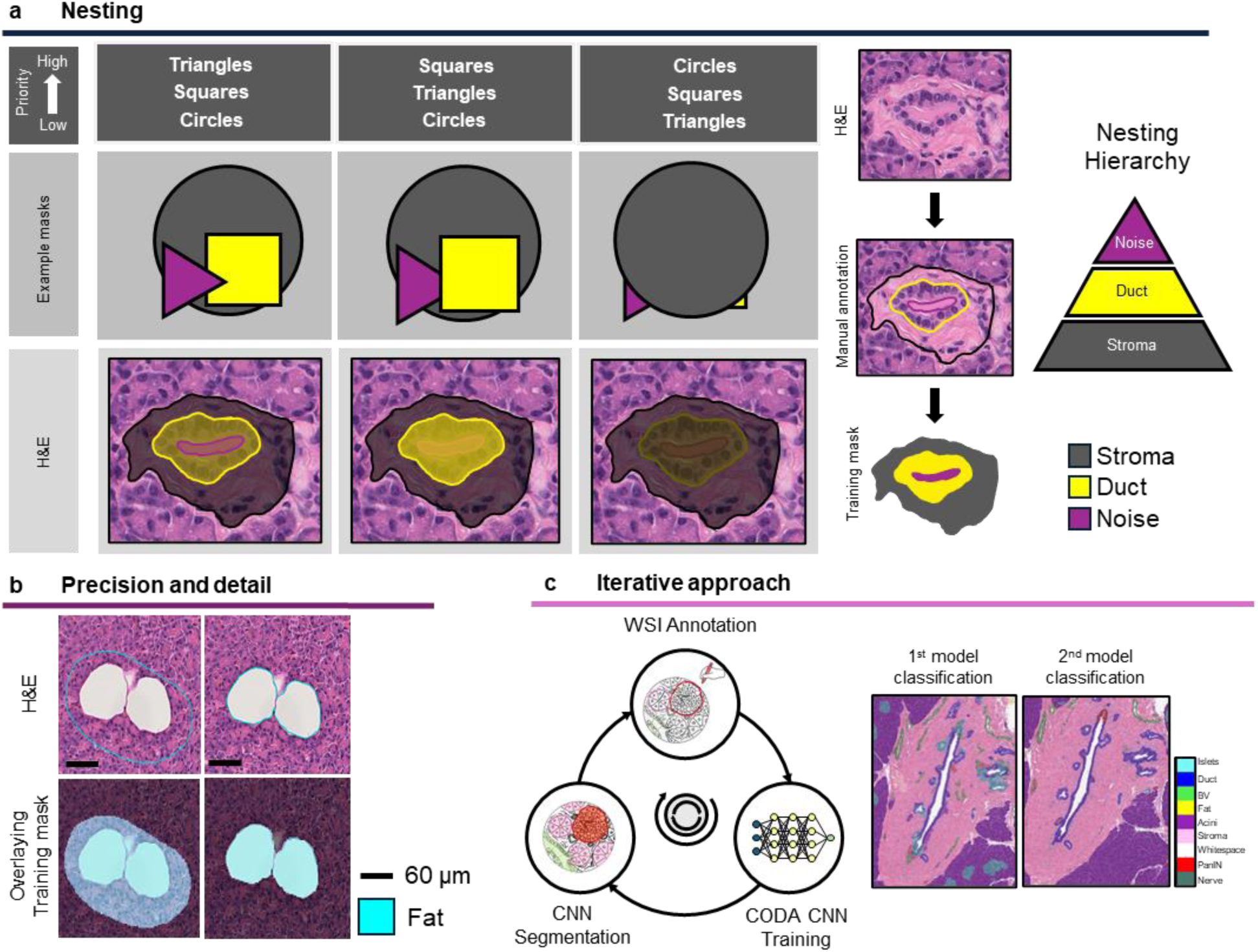
Annotation hierarchy and quality control for CODA deep learning model training. a. Hierarchical nesting diagram illustrating tissue classification levels. b. Comparison of adipocyte annotation precision within pancreatic acinar tissue: suboptimal classification (left) versus optimal annotation (right). c. Iterative annotation workflow demonstrating the targeted addition of annotations in misclassified regions to enhance model performance.

**Figure 4:**
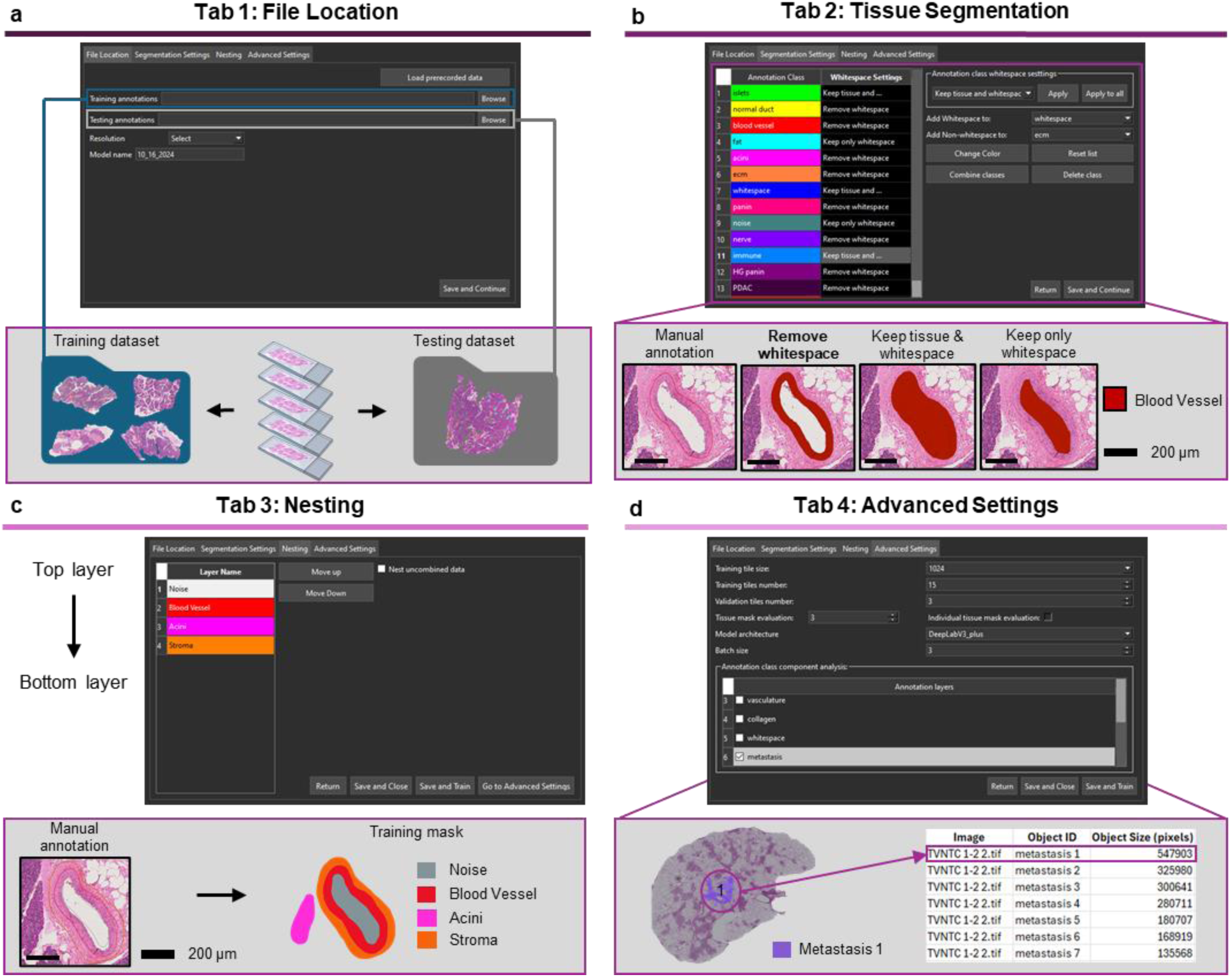
CODAvision graphical user interface (GUI) tabs. a. File Location tab: Configuration of dataset paths, model name, and training image resolution. b. Segmentation Settings tab: Parametrization of whitespace management c. Nesting tab: Configuration of nested annotation hierarchy for overlapping tissue annotations. d. Advanced Settings tab: Modification of CNN hyperparameters and selection of annotation classes for component analysis.

**Figure 5:**
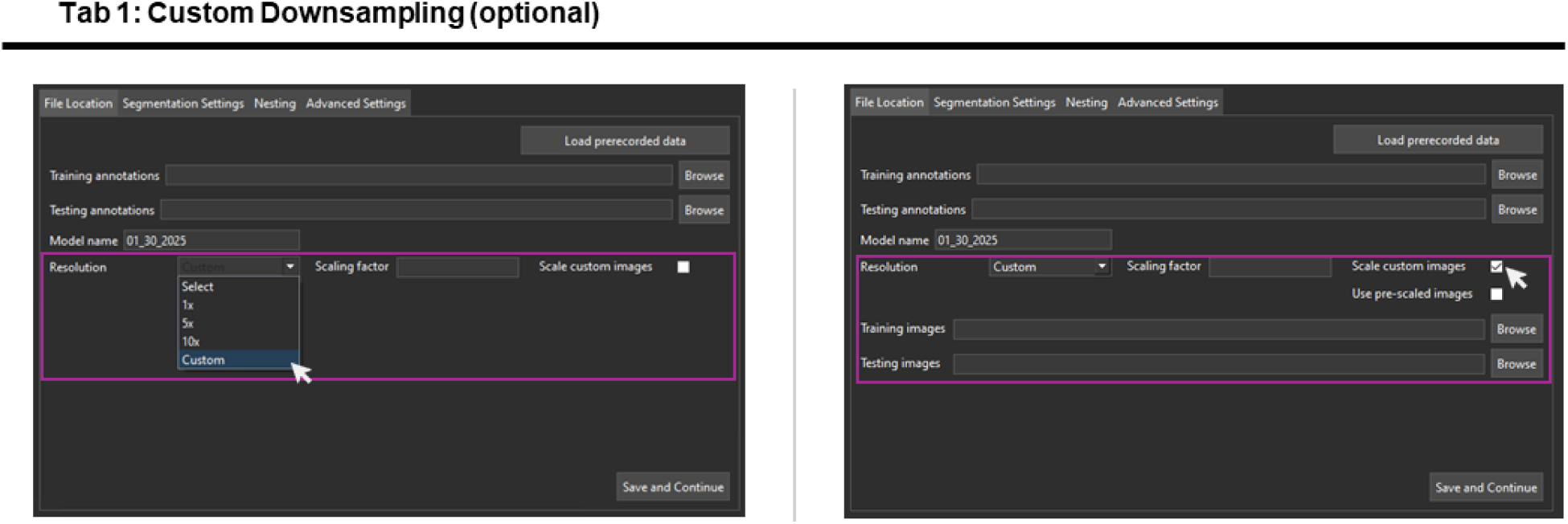
Optional tab for custom downsampling of images or for providing pre-downsampled files to the GUI.

**Figure 6:**
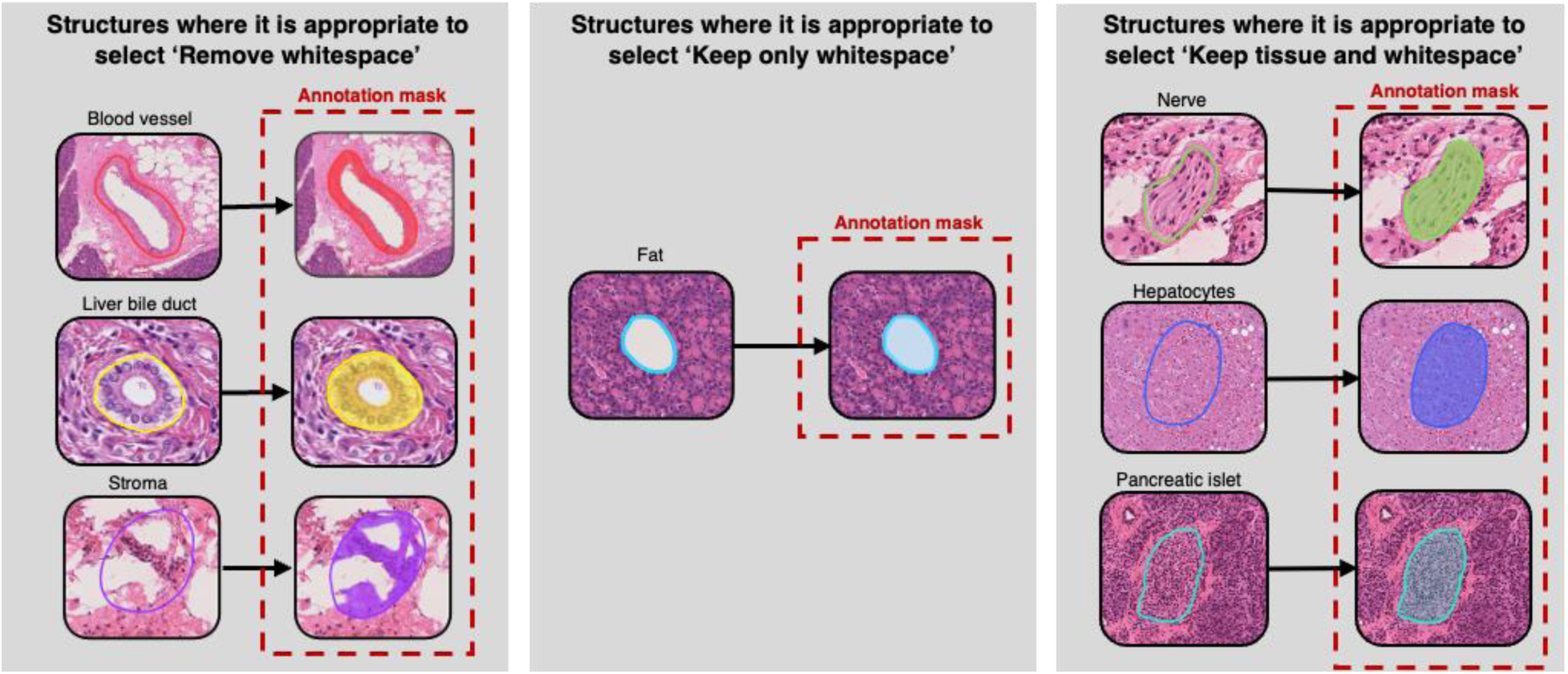
Example annotations where each whitespace management option is best.

### Procedure 3: Image pre-processing and model training

#### Tissue mask thresholding

##### • Timing: 5-10 min

1. Upon completing the parameter configuration in the CODAvision GUI, the software will initiate execution and begin downsampling the training images to the resolution specified in Procedure 2, Step 6.
2. Next, a popup window will appear, enabling interactive selection of a threshold value to separate tissue pixels from background pixels. (**Fig 7**)
3. In the popup window, an image will be displayed. Double click on a region of the image containing both tissue and whitespace.
4. The popup window will reload and display a magnified view centered on the selected region. Confirm the selection or choose a different region until a suitable area with sufficient tissue and background is identified.
5. nce the region is confirmed, a new window will prompt the user to adjust the threshold cutoff until the background is detected optimally:
6. After selecting the desired threshold, another full-size image will load to repeat the process until the desired number of images has been assessed. The average threshold value from this process will then be applied to all images in the cohort. CRITICAL: Tissue mask threshold optimization is required only once. If retraining the model on the same dataset, a popup window will prompt the user to either ‘Keep current tissue mask evaluation’, which reuses the previously determined threshold and skips this step, or ‘Evaluate tissue mask again’, which initiates a new round of threshold selection. If the ‘Individual tissue mask evaluation’ checkbox (see Procedure 2, Step 25) was selected and the user chooses to ‘Evaluate tissue mask again’, a prompt will appear allowing manual selection of specific tissue masks that the user desires to remake. These tissue masks are located in the training annotation path in the downsampled images folder. Alternatively, selecting ‘Redo all images’ will apply the thresholding process to the entire dataset. <u>CODAvision image processing and model training (</u>**<u>Fig 8</u>**) **• Timing: 2 h**
7. Following successful completion of tissue mask optimization, the image pre-processing and model training will proceed automatically. CRITICAL: This step of the protocol is completed automatically by the GUI once the training is initiated in Procedure 2. The user has no direct steps in this section, but the steps followed by the code are described below for clarity.
8. While executing, the code will output several text statements to the command window, in the following order:

a. The model name.
b. A message indicating the location where the model metadata will be saved.
c. For each annotated image, a message will appear indicating the status of the annotation metadata import:

i. If the annotation data were loaded in a previous training and the annotations and model parameters have not changed, the message will state that the metadata for this image was previously loaded.
ii. If the annotation data are being loaded for the first time, the following checkpoint messages will be displayed: (1) Confirmation that the .xml annotation file has been converted to.pkl format; (2) Notification that an annotation mask has been generated and saved as a .png file; (3) Confirmation that smaller image patches have been cropped around each annotated region using the annotations mask, producing ‘bounding boxes’ of the RGB image and mask for downstream processing.
9. After loading all annotation metadata, the code will output the total size of the training dataset, including the percentage of the training dataset contributed by each class. CRITICAL: It is recommended to have a well-balanced dataset with many examples of each class. The codes will automatically calculate the percentage of annotations for each class. We suggest that the minimum class will make up at least 5% of the total annotations must represent each class to ensure sufficient heterogeneity is provided for each annotation class and that the heterogeneity of the annotation classes is represented roughly equally. If any class if under the 5% threshold, we recommend the user attempt to add manual annotations of the least prevalent class through further annotation of the training images or through addition of new training images.
10. After the metadata is loaded for all annotation images and the composition of the training dataset is calculated, training and validation tiles will be constructed. **How to generate robust deep learning models Box 6** Construction of a large and diverse training dataset is important to generate robust deep learning models. We augment the manual annotations in several ways to increase the heterogeneity of the training data. For each training tile, a large zero-value image of size 10,000 × 10,000 × 3 pixels is generated. Bounding boxes containing processed manual annotations are randomly added to the tile until the tile is >55% filled. Each iteration, the least represented class in the tile is determined, and an annotation bounding box containing that class is added to the tile. Approximately 50% of tiles are augmented by adjusting the hue (scaling RGB channels independently within a range of 0.88 to 1.12), scaling (within a range of 0.6 to 0.95 for downscaling and 1.1 to 1.4 for upscaling), rotating (at random angles from 0 to 355 degrees in increments of 5 degrees), and applying a Gaussian blur (with sigma values randomly selected from a predefined set, e.g., 1.05, 1.1, 1.15, 1.2). Each bounding box is thus added many times to the training tiles, but each time in a new position surrounded by different neighbors, and augmented in several ways. Once >55% of the tiles are filled, these large tiles will be cropped to a size of 1024 × 1024 × 3 pixels (or the custom size if defined by the user in the advanced settings tab of the GUI) and saved.
11. After the predefined number of training and validation images have been created, model training will start. All training and validation tiles will be normalized through zero-centering. The model will be trained for a maximum of 8 to 10 epochs depending on the CNN chosen (see Procedure 2, Step 25) and validated three times per epoch. Model training stops according to an early stopping patience of 6.
12. Following model training, annotation masks will be generated for the test dataset as detailed in Step 7 of this Procedure.
13. The testing image(s) will be segmented, with the resulting segmentation masks and colorized overlays saved as .png and .jpg files, respectively. This output will be compared to the metadata collected in Step 11 of this Procedure to generate a confusion matrix.
14. The images specified in the training images folder will be segmented, with the resulting segmentation masks and colorized overlays saved as .png and .jpg files, respectively.
15. The bulk tissue composition of the images in the training images folder will be calculated and saved in a .csv file.
16. (Optional) Component analysis will be calculated for selected annotation classes in the ‘Advanced settings’ tab described in Procedure 2 and saved in a .csv file. This step would be skipped if no classes were selected.

#### Generated outputs for Procedure 3

Model hyperparameters, trained weights, and results will be stored in the training path file location. A summary of input and output file formats for the CODAvision workflow is given in **Fig 9**. This workflow generates outputs per annotated image, per trained model, and per segmented image.

1. ’data_py’ folder:

a. Subfolder for each training image containing:

i. ‘annotation.pkl’: processed vertex coordinates corresponding to the manual annotation data saved in each .xml file
ii. ‘view_annotations.png’: grayscale mask images containing pixel labels for each annotated region, considering the whitespace management and the nesting order.
iii. ‘model_name_boundbox’ subfolder: contains cropped RGB and mask images corresponding to each annotated region identified in ‘view_annotations.png.’

1. ‘im’ subfolder: contains the RGB copy of the bounding boxes.
2. ‘label’ subfolder: contains the corresponding mask of the bounding boxes. CRITICAL: The ‘data_py’ folder contains metadata used internally for image processing and training tile generation, not for analysis.
2. ’check_annotations’ folder: contains colorized masks of the training images with annotation patches highlighted in colors defined in Tab 2 of the GUI. CRITICAL: Review the ‘check_annotation’ images to verify that your annotations are correctly overlaid on the downsampled images. Pay close attention to the whitespace handling (adjust the whitespace settings in Tab 2 of the GUI if needed, or regenerate the tissue mask images). Also, ensure the nesting order is correct; if necessary, revise the annotations or modify the nesting settings in Tab 3 of the GUI.

a. ‘Model_name’ folder
b. ‘training’ folder: contains image tiles used for model training
c. ‘validation’ folder: contains image tiles used for model validation
d. ‘best_model_X.keras’: weights of the CNN for the best training epoch, where X is the name of the model architecture that was trained.
e. ‘X.keras’: weights of the trained CNN, where X is the name of the model architecture that was trained.
f. ‘net.pkl’: annotation parameterization settings.
g. ‘model_color_legend.jpg’: color map used in colorized segmented images.
h. ‘model_evaluation_report.pdf’: detailed performance report. (**Fig 10**)
i. ‘confusion_matrix_X.png’: confusion matrix with precision, recall, and accuracy metrics, where X is the name of the model architecture that was trained. CRITICAL: We suggest a minimum overall accuracy score greater than 90%, and a minimum per-class precision and recall exceeding 85% for acceptable training results. For models that do not meet these standards, we suggest users add additional training annotations and retrain to improve the metrics.
3. Downsampled image folder named based on the chosen resolution (e.g., ‘1x’, ‘5x’, ‘10x’, or a user defined custom scale)

a. ’TA’ subfolder: ‘tissue area’ masks for the training images.
b. ’classification_model_name’ subfolder, where ‘model_name’ is the user-defined name of the trained model. This subfolder will contain:

i. grayscale segmentation masks of each training image in .tif format

1. ‘check_classification’ subfolder: colorized segmentation masks in .jpg format.
ii. ‘image_quantifications.csv’: pixel count and tissue composition for each training image.
iii. ‘annotation_class_name_count_analysis.csv’: object count and size analysis for each class. CRITICAL: To visually assess the model accuracy, review the ‘check_classification’ images and search for areas that are misclassified.

**Figure 7:**
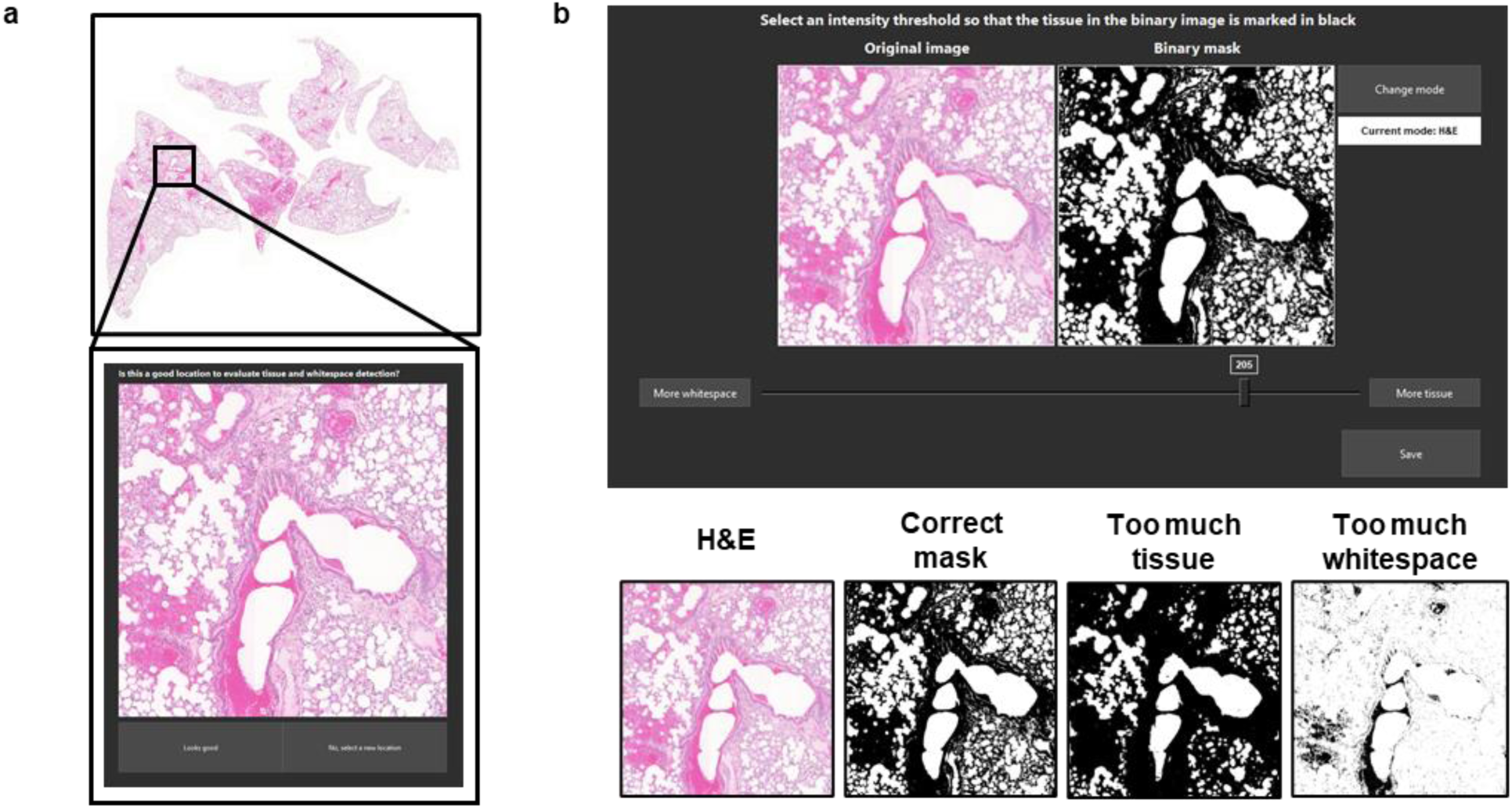
Tissue threshold selection interface. a. Region of interest selection in a lung whole-slide image (WSI) to optimize threshold values. b. Threshold adjustment interface with optimal tissue-background discrimination demonstrated in the lower panel.

**Figure 8:**
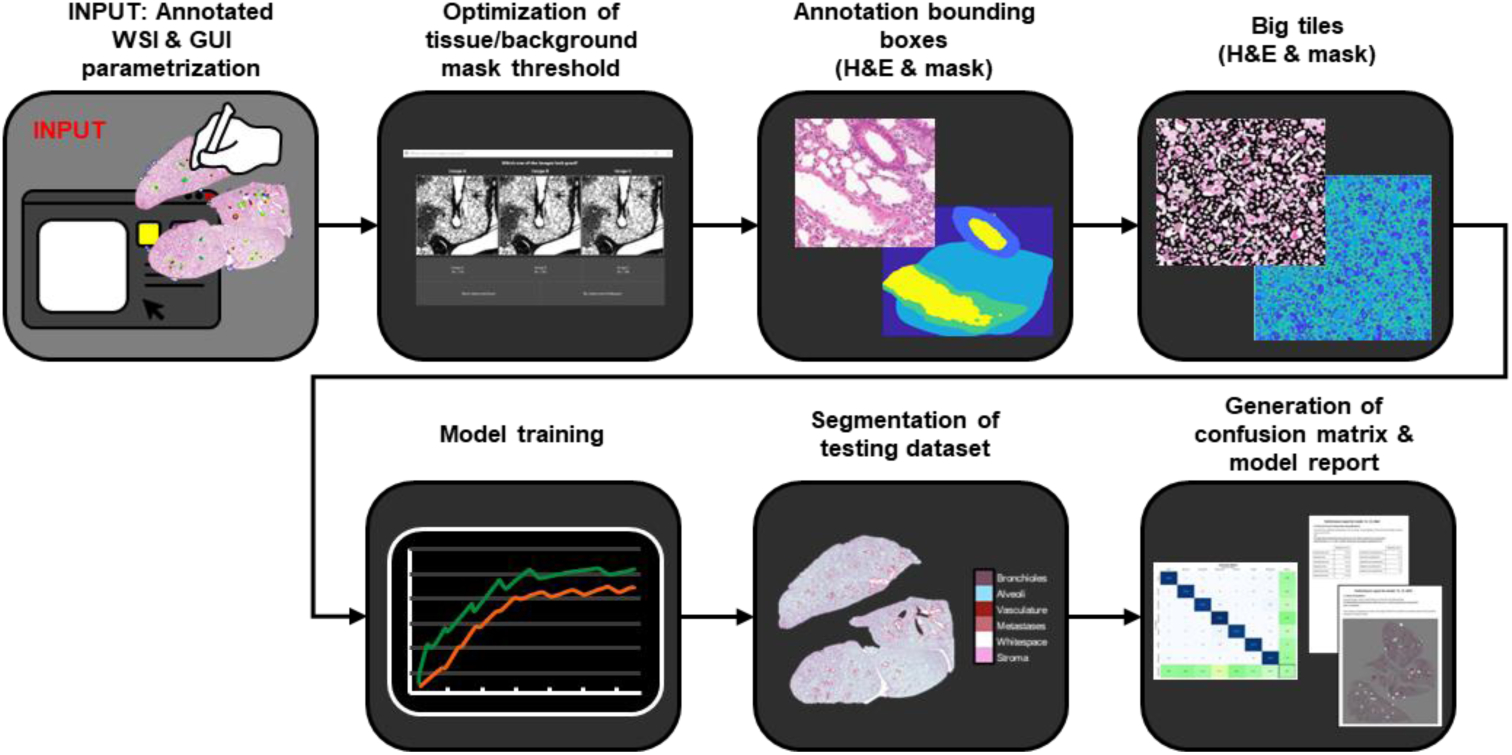
Visualized workflow of Procedure 3.

**Figure 9:**
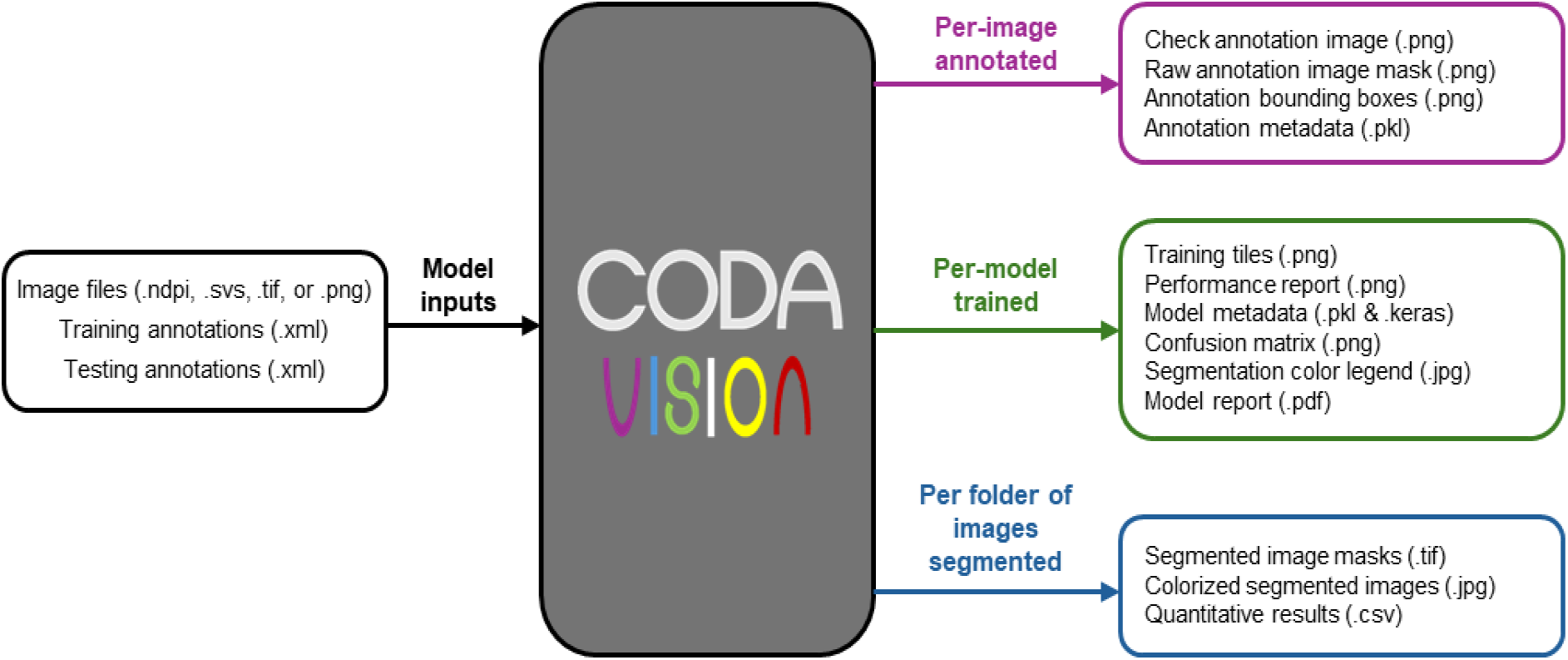
Overview of CODAvision inputs and output data formats.

### Procedure 4: Model optimization

#### Timing: 1 – 10 h

If a trained model is of unsatisfactory quality (overall accuracy <90%, minimum precision or recall <85%, or visually poor performance), follow this procedure to retrain the model efficiently. It is common when starting a new project to train 2 – 3 models in quick succession before achieving satisfactory results. We describe here four techniques to improve model performance (**Fig 11**).

1. First, determine if there is human error in the GUI settings. If yes, reconfigure the settings and retrain the model without adding new annotations.

a. Review the color overlay images inside the ‘check_annotation’ folder to determine if the annotations contain errors or model parameters were defined incorrectly.

i. Does the whitespace management look correct? For example, in an annotated blood vessel, are the background pixels inside the lumen correctly extracted as whitespace? If incorrect, consider adjusting the whitespace management in tab 2 of the GUI (see again **Fig 6** for examples).
ii. Does the nesting order in the images look correct? For example, if noise was annotated inside of a duct, is that region defined in the check_annotation file as noise or as duct? If incorrect, consider adjusting the nesting order in tab 3 of the GUI (see again **Fig 3** for examples).
iii. If the nesting order and whitespace management look correct, is anything else noticeable in the file? Are any structures incorrectly annotated? Consult the supplementary annotation guide (**supplementary file 1**) for detailed best practices in annotation and correct these errors by editing the annotations where necessary. CRITICAL: Biological images contain complex and subtle structures. Human error in manually annotating images is common especially for new users. If errors are identified in the annotated files, correct them and retrain, and remember to also correct for errors in the testing dataset.
b. the tissue area masks to determine whether the threshold is correctly separating the tissue and background (see again **Fig 7** for examples). If this threshold looks incorrect, delete the subfolder named ‘TA’ so that a new threshold may be calculated upon retraining.
2. Second, view the images saved inside the folder named ‘check_classification’ to visually assess the model performance and identify misclassified regions. These files show a color overlay of the model segmentation across whole images.

a. Review several images and identify common ‘patterns’ in misclassification. For example, stroma consistently misclassified as vasculature, or background pixels consistently misclassified as fat.
b. Determine whether there are any structures present in the images that were missed during the previous model training and were not included as an annotation layer (for example, if blood vessels were not included as a label but are present in the images). If identified, determine whether this structure should be combined with a current annotation layer or added as an additional annotation layer.
3. After correcting any problems caused by human error in model parametrization and retraining the model, next the user may improve the model results through further annotation. Focusing on the list of patterns in misclassification generated in Step 2 of this Procedure, add annotations to the identified problem areas. CRITICAL: If the model frequently misclassifies unseen structures when applied to images beyond the training set, consider augmenting the training dataset with entirely new training images, rather than adding additional annotations to images previously used for training.
4. If all the human errors have been corrected and the training dataset is extensive, the user may also consider tweaking model parameters in the advanced settings tab of the GUI. CRITICAL: When retraining a model using identical annotations (e.g. same training and validation tiles) but with a different architecture, load previous model settings using the ‘Load prerecorded data’ button and modify the model architecture via the advanced settings tab in the CODAvision GUI (Procedure 2, step 25). This approach eliminates the need to repeat steps 1-10 of Procedure 3, allowing rapid retraining of models.
5. Repeat the steps of procedures 1 – 4 to retrain the model and reoptimize the annotations and model parametrization until satisfactory results are obtained. When retraining a model, consider using the ‘Load prerecorded data’ option to prepopulate the model parameters.

**Figure 10:**
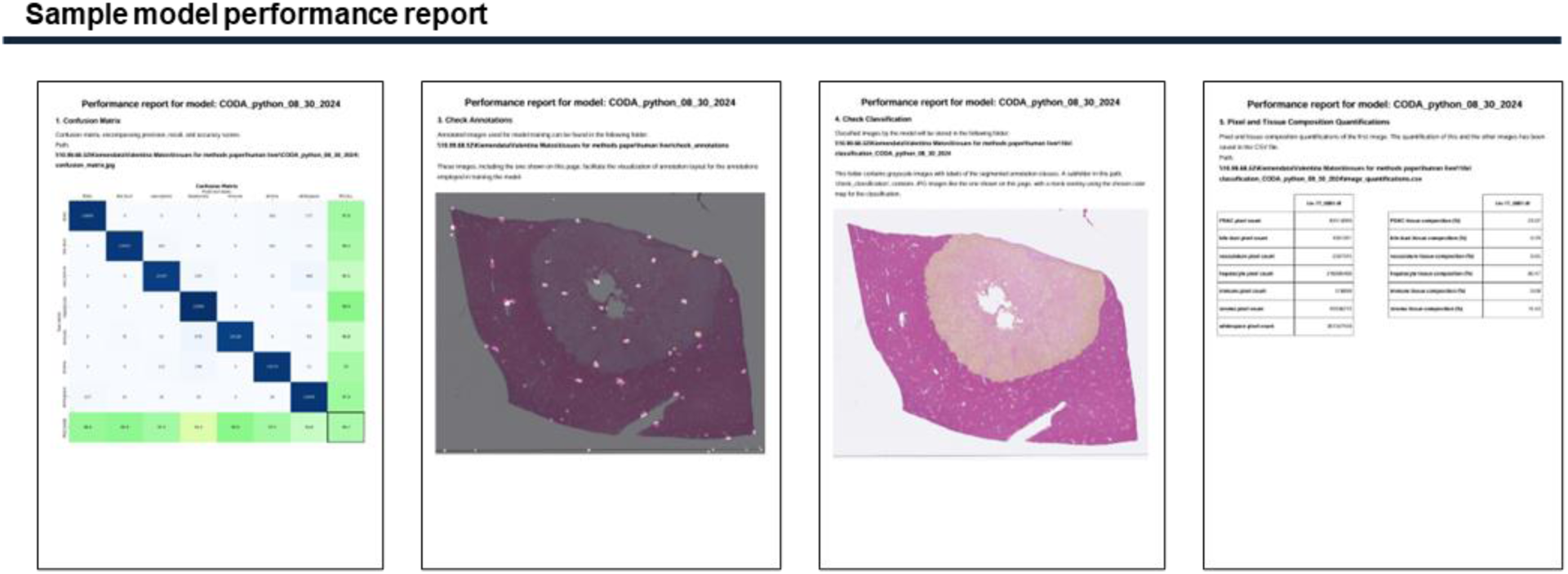
Sample PDF report summarizing the performance metrics and training parameters of a model trained to recognize normal hepatic cells and cancer in human liver histology.

**Figure 11:**
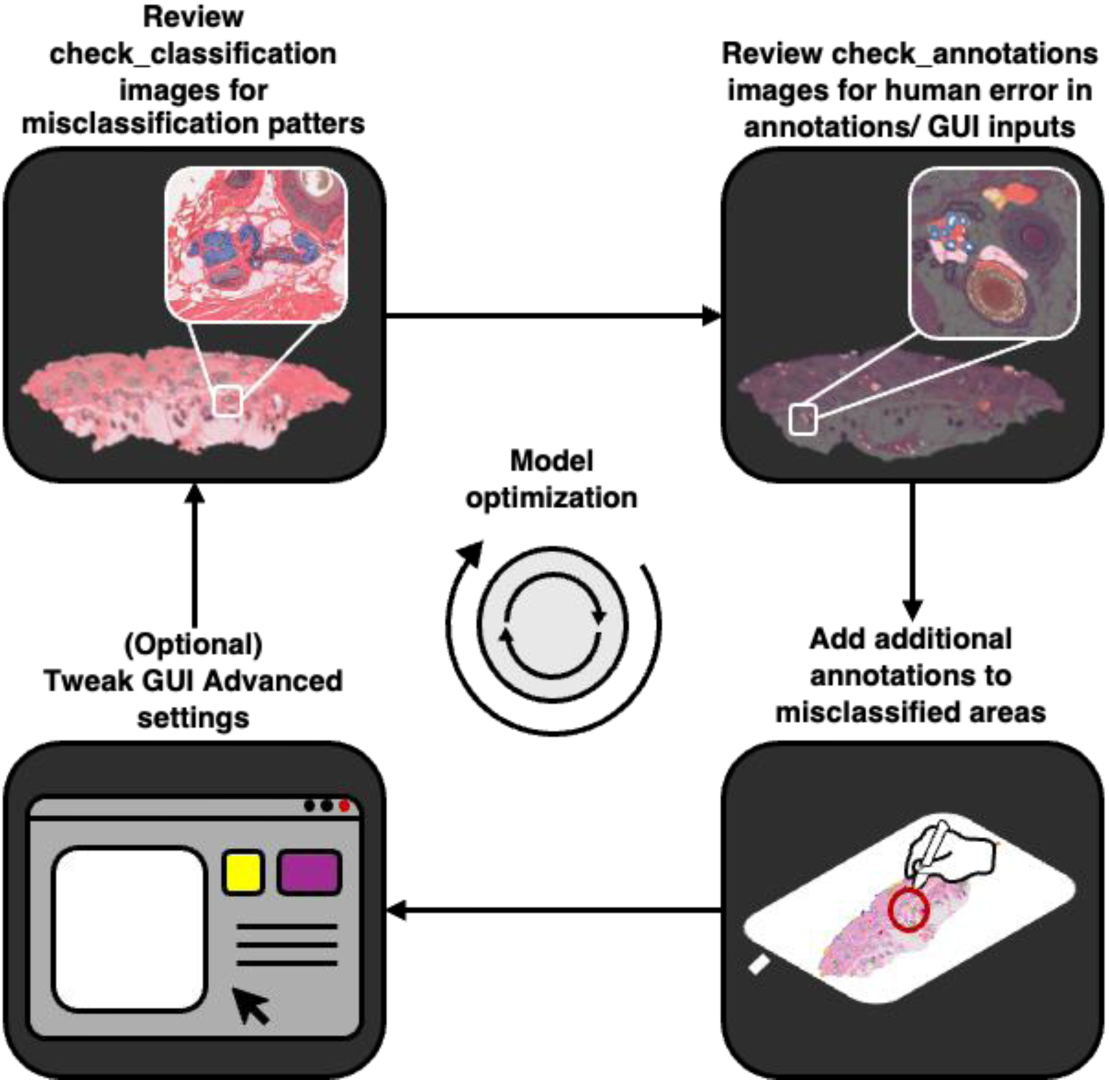
Suggested solutions to incrementally improve model performance.

### Procedure 5: Image segmentation using a pretrained model

#### Timing: 10 – 15 min

After training a model, the user may segment additional images beyond the original training set.

CRITICAL: To ensure high model performance, only images with the same resolution as the training data set should be classified with a pretrained model.

1. Run the CODAvision.py code to restart the CODAvision GUI.
2. Load the training weights of the model you wish to use (**Fig 12A**). To do this, either select ‘Load prerecorded data’ and select the folder containing the ‘net.pkl’ file you wish to use, or manually input text in the ‘Training annotations’, ‘Resolution’ and ‘Model name’ fields. When these fields are completed and a trained model is detected, a green ‘Classify images’ button will appear. Click this button to proceed to the ‘Segment images’ tab.
3. After proceeding to the ‘Segment images’ tab (**Fig 12B**), input folders containing images you wish to classify. Click the ‘Browse’ button to add a desired folder to the table list and repeat until all desired folders are added.
4. (Optional) Modify the segmentation color map. Change the colors of annotation classes in the table on the left. This new color overlay may be visualized in real time to ensure readability through application to the sample image displayed in the center of the tab. The updated color map will be used to generate the colorized segmented images. CRITICAL: If the objective is only to modify the color map of previously classified images, input the root directory path of images that have been classified by the model specified in step 2.a, and click ‘Apply’. The code will begin execution, bypassing the segmentation and will change the colorized segmented images.
5. (Optional) Perform annotation class component analysis. Select the annotation classes to quantify in the table on the right.
6. Click ‘Apply’ to segment all new images in all folders added to the segmentation table and to perform the quantitative analyses as defined.

**Figure 12:**
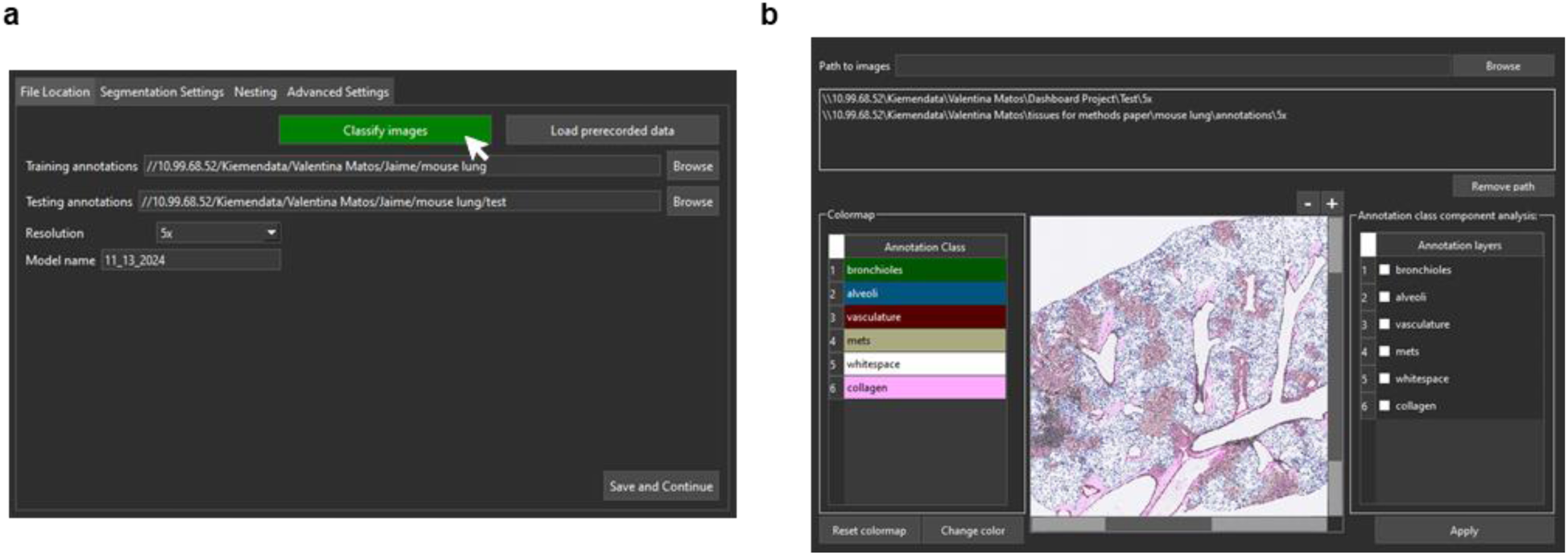
GUI tab for classifying additional images with pretrained models. a. File location tab showing the ‘Classify Images’ button, which appears when a trained network is detected in the specified data folder. b. Interface for selecting new image set paths to classify with the pretrained model, modifying segmentation mask color map, and performing additional annotation class component analysis.

#### Timing

The estimated processing times provided below are based on the training of sample Dataset 1 (human liver histology), trained at the selected magnification of 10× (1 µm / pixel resolution), performed using a workstation equipped with an NVIDIA GeForce RTX 4090 GPU running Windows 11. These estimates reflect the processing time for an experienced CODAvision user using a GPU equipped computer. Processing times will increase substantially if the software is run without GPU support. Note that users following the CODAvision workflow for the first time may require additional time to familiarize themselves with the annotation best practices and GUI guided parametrization.

- Procedure 1: Building annotation dataset

- Steps 1-12: 8-10 h
- Procedure 2: GUI guided parametrization

- Steps 1-26: 5 min
- Procedure 3: Image pre-processing and model training

- Steps 1-5, tissue mask thresholding: 5-10 min
- Steps 6-16, CODAvision image processing and model training: 2-3 h
- Procedure 4: Testing a trained model and generating a performance report

- Steps 1-4: 5 min
- Procedure 5: Model optimization

- Steps 1-5: 1-10 h
- Procedure 6: Segmenting an image with a pretrained model

- Steps 1-9: 2-3 min

### Troubleshooting

| Step | Problem | Possible reasons | Solution |
| --- | --- | --- | --- |
| **Procedure 2** |  |  |  |
| 1 | Unable to install cudatoolkit and cudnn dependencies | SSL certificate verification issues | Make sure that OpenSSL is installed and properly configured on your system. You can download and install OpenSSL from https://slproweb.com/products/Win32OpenSSL.html. After downloading OpenSSL add its bin directory to the environment variables |
| 1 | Unable to download CODAvision | Git not installed on computer | Download git from the following website (https://git-scm.com/downloads/win) and restart your computer |
| 2 | Cannot execute CODAvision.py | Dependencies not properly installed | Ensure .dll path for OpenSlide is located in environment paths |
| **Procedure 3** |  |  |  |
| 7 | Model training failure | Existing model with same name in file directory | Rename the model or delete the previous model's folder |
| 8 | Code cannot identify annotation .xml file for training mask | Filename mismatch between annotated images, .xml files, and downsampled .tif images | Ensure all filenames are consistent and rerun the code |
| 11 | GPU out of memory | Model architecture or batch size too resource-intensive | Choose a lighter model architecture or reduce batch size to match computer specifications |
| 11 | Validation loss increases after a certain number of epochs (overfitting) | Insufficient or incorrectly annotated training data | Increase number of annotated images or improve annotations |
| 12 | Model testing failure | Testing annotations lack classes present in training dataset | Include annotations for all classes in the training dataset |
| 12 | Confusion matrix shows all pixels classified into a single class | Numerical instability during GPU training | Retrain and test the model using CPU processing instead |
| **Procedure 4** |  |  |  |
| 1,2 | Confusion matrix shows low precision and recall for light contrast classes similar to background | Poor tissue background cut-off | Repeat CODAvision workflow with different tissue background cutoff value |
| 2 | Misclassification in images outside training dataset | Insufficient annotation class diversity or heterogeneity | Include more annotation classes for morphologically different structures, create more heterogeneous annotations, or increase training dataset with additional annotated images (refer to supplemental annotation guide in **supplementary file 1**) |

### Anticipated results

This protocol provides detailed steps to build highly customizable semantic segmentation models. We provide comprehensive guidelines to (1) determine the optimal number of target labels for a dataset given the scientific question, and (2) annotation best-practices to create effective training and testing datasets. We also provide instructions to operate our user interface, including streamlining the processes of parameter configuration, model architecture selection, and model training.

After successfully completing the protocol, the user will have generated several files in user-friendly formats. This protocol enables users to rapidly build highly customizable segmentation models for biological research. In this section, we provide use-cases to demonstrate the value of the CODAvision outputs across several research areas.

To demonstrate the versatility of CODAvision, we applied the workflow to four distinct research tasks (**Fig 13**). For each application, we trained using a DeepLabv3+ ResNet50 model architecture. Below, we briefly describe each use case and the analyses enabled by CODAvision.

**Figure 13:**
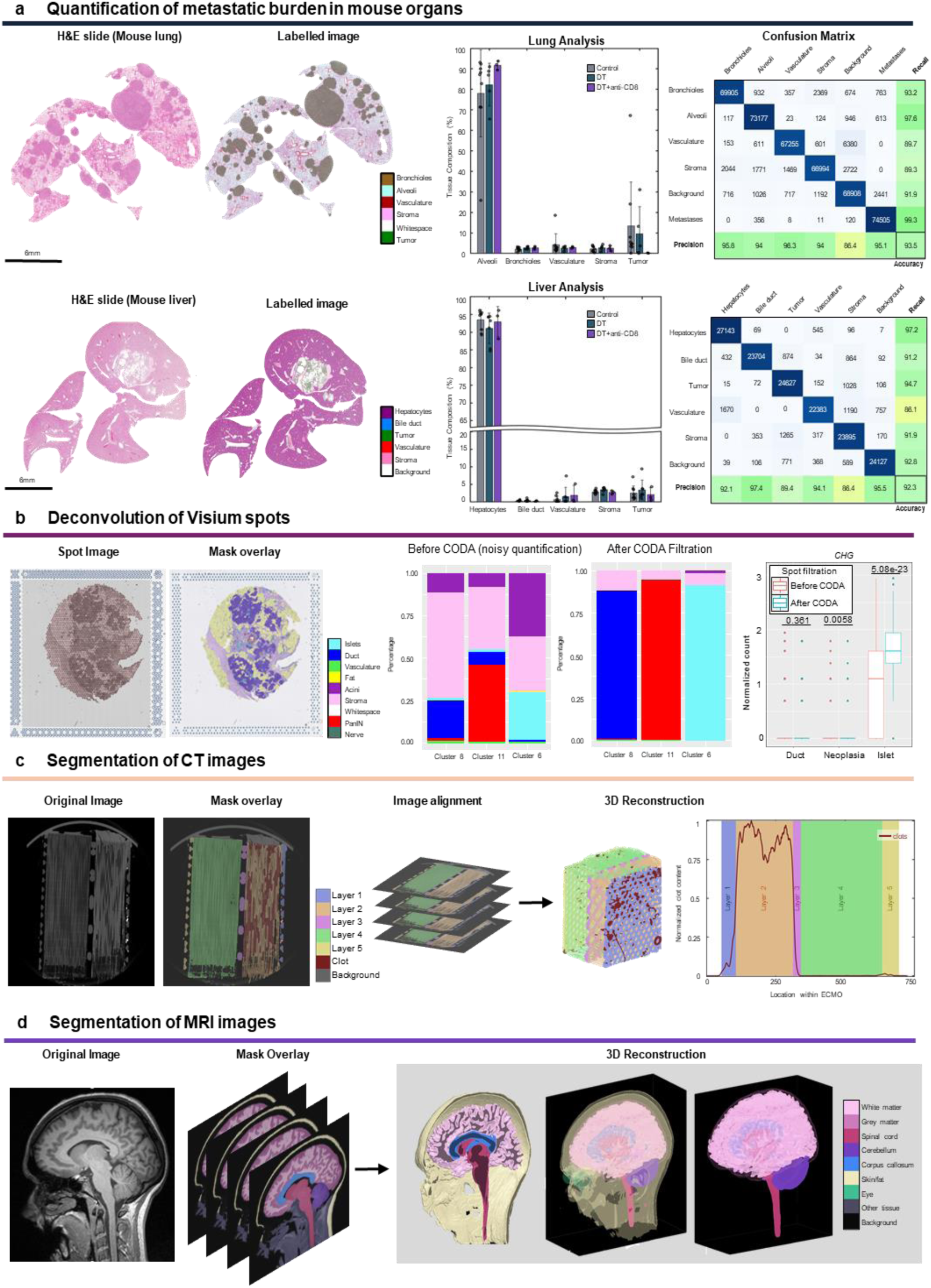
Example applications of CODAvision to different medical image types. a. Quantification of metastatic burden in mouse lung and liver through semantic segmentation and automated tissue quantification. b. Deconvolution of Visium spots using CODAvision segmentation to reduce noise and improve quantification accuracy within each cluster. c. Segmentation of CT images. By leveraging different annotated labels, structures with similar gray intensity values were successfully separated into distinct anatomical regions. d. Segmentation of MRI images enables visualization of the anatomy of the brain and spinal cord.

First, we quantified microanatomical structures in liver and lung histology from a syngeneic mouse model of pancreatic cancer (**Fig 13A**). We demonstrate that CODAvision can be used to quantify metastatic burden and organ composition. This histological dataset was generated through an *in vivo* experiment originally described in the work cited.^51^ Here, we trained separate models for each organ. We segmented six structures in the mouse lungs and seven structures in the mouse liver. In both models we achieved >90% overall accuracy and >85% per-class precision and recall. In so doing, we were able to demonstrate that in this mouse model, the lungs metastases grow larger more quickly, are more numerous, and are more solid than the liver metastases. The pipeline simultaneously analyzed the composition of other key tissues (bronchioles, alveoli, stroma, and vasculature in the lungs, and hepatocytes, bile duct, stroma, fat, and vasculature in the liver), demonstrating its utility for comprehensive tissue composition analysis in preclinical models.

Second, we segmented cell types in human pancreas histology to deconvolute spot-based spatial transcriptomic data (**Fig 13B**). Deconvolution of spatial transcriptomics data has emerged as a vital process for honing in on gene expression signatures of target cells that may make up a small fraction of the sampled tissue.^31,32,52–54^ This dataset, consisting of an H&E image and 10× Genomics Visium spatial transcriptomics outputs, and the computational method for deconvolution using segmentation results, were originally described in the work cited.^32^ Here, we segmented nine structures of pancreatic microanatomy, including normal pancreatic ducts and pancreatic precancers, achieving >90% overall accuracy and >80% per-class precision and recall. The cell type labels at the coordinates of each 55µm radius Visium spot were extracted and used to assess the cellular purity of each spot. This deconvolution allowed us to clearly determine the gene expression of pancreatic precancerous cells and to eliminate confounding gene expression signatures contributed by non-neoplastic cells within the same Visium spot.

Third, to demonstrate CODAvision’s broad applicability to biological images beyond histology, we segmented functional components of ECMO membranes in CT (**Fig 13C**). ECMO is a device used in critical care medicine to provide blood oxygenation for patients suffering from conditions that impact lung function such as COVID-19.^55^ The ECMO membrane is made up of 5 distinct fibrous layers contained within either a square or cylindrical exterior casing. Blood is perfused from the patient and through the multiple matrix design layers and the hollow fiber layers of the ECMO where gas exchange occurs before the blood is returned to the patient.^56^ The dataset shown here was originally presented in the cited work^57^ where CT images of used ECMO membranes were analyzed to determine which of the five fibrous layers contained the highest composition of blood clots (a common cause of ECMO failure). Due to the highly specialized nature of the desired segmentation, more common methods for CT segmentation such as quantification of changes in density need further optimization. CODAvision can be used to automate segmentation across every CT slice, ensuring precise and consistent identification of layers and clot formations. We therefore trained a segmentation model to detect seven structures in the ECMO device including the five fibrous layers and blood clots, achieving >90% overall accuracy and >80% per-class precision and recall. After training, we applied the segmentation model to classify the 985 serial images making up the 3D CT data. Using the outputted .tif segmentation masks, we constructed a project-specific analysis to quantify the composition of blood clot in each of the fibrous layers of the ECMO, identifying layer 2 as a region with substantially higher percentage of clots. This analysis, possible only through segmentation of the subtle textures that define the distinct ECMO layers, will enable future design of more efficient ECMO devices that are less susceptible to blockage by blood clots.

Finally, to further highlight the versatility of CODAvision, we applied our method to an MRI dataset of the human brain (**Fig 13D**). Here, we desired to rapidly construct a highly specific model to distinguish the anatomical components of the brain including white matter, grey matter, the cerebellum, and non-brain structures including the human eyes. We annotated these structures and trained a segmentation model with CODAvision, achieving >90% overall accuracy and >80% per-class precision and recall. After model training, we applied the segmentation model to the 157 serial images making up the 3D MRI data. Using the cited method^25^ we transformed the segmented .tif images outputted by CODAvision into a 3D printable .stl file. This enabled us to 3D print the anatomy of the brain.

## Data availability

Sample datasets are available at the following link: https://drive.google.com/drive/folders/1K-wY_ArVGbEhebQD4AjOeERwx6-4Fw3G

## Code availability

The CODAvision package is available in the following repository: https://github.com/Kiemen-Lab/CODAvision.

## Author contributions statements

ALK, DW, and VM-R conceived the project. VM-R led the computational analysis, with help from JG-B, AF, LD, TN, SJ, AS, YS, SN, EH, PN, JS-HW, EL-O, ATFB, LDW, LK, EF, JC, CS, D-FD, MPM, JJS, OJTM, JOL, AR, RHH, and AM-B provided sample datasets and guidance. VM-R and ALK drafted the manuscript, and VM-R drafted the protocol and supplementary guides, which all authors edited and approved.

## Acknowledgments

We would like to thank the following sources of support: U54CA268083; Lustgarten Foundation-AACR Career development award for pancreatic cancer research in honor of Ruth Bader Ginsburg; Susan Wojcicki and Denis Troper; The Carl and Carol Nale Fund for Pancreatic Cancer Research; the Rolfe Pancreatic Cancer Foundation; Fight Cancer Stay Positive; The Sol Goldman Pancreatic Cancer Research Center; The Johns Hopkins University Data Science and Artificial Intelligence Program. National Institutes of Health (NIH) Grant P51 OD011092. This work was partially supported by the Ministerio de Ciencia, Innovación y Universidades, Agencia Estatal de Investigación, MCIN/AEI/10.13039/501100011033, under grant PID2023-152631OB-I00, co-financed by European Regional Development Fund (ERDF).

## Competing interests

The authors declare no competing interests.

## Notes

https://github.com/Kiemen-Lab/CODAvision

### Competing Interest Statement

The authors have declared no competing interest.

## References

1 Chan, H. P., Samala, R. K., Hadjiiski, L. M. & Zhou, C. Deep Learning in Medical Image Analysis. Adv Exp Med Biol 1213, 3–21 (2020). 10.1007/978-3-030-33128-3_1

2 Lundervold, A. S. & Lundervold, A. An overview of deep learning in medical imaging focusing on MRI. Z Med Phys 29, 102–127 (2019). 10.1016/j.zemedi.2018.11.002

3 Salto-Tellez, M., Maxwell, P. & Hamilton, P. Artificial intelligence-the third revolution in pathology. Histopathology 74, 372–376 (2019). 10.1111/his.13760

4 Bera, K., Schalper, K. A., Rimm, D. L., Velcheti, V. & Madabhushi, A. Artificial intelligence in digital pathology - new tools for diagnosis and precision oncology. Nat Rev Clin Oncol 16, 703–715 (2019). 10.1038/s41571-019-0252-y

5. Paul, J., et al. Digital transformation: A multidisciplinary perspective and future research agenda. Int J Consum Stud 48 (2024). ARTN e13015 10.1111/ijcs.13015

6 Echle, A. et al. Deep learning in cancer pathology: a new generation of clinical biomarkers. Brit J Cancer 124, 686–696 (2021). 10.1038/s41416-020-01122-x

7 Ghahremani, P. et al. Deep learning-inferred multiplex immunofluorescence for immunohistochemical image quantification. Nat Mach Intell 4, 401-+ (2022). 10.1038/s42256-022-00471-x

8 Zhang, D. W. et al. Inferring super-resolution tissue architecture by integrating spatial transcriptomics with histology. Nature Biotechnology 42 (2024). 10.1038/s41587-023-02019-9

9 Lu, M. Y. et al. Data-efficient and weakly supervised computational pathology on whole-slide images. Nat Biomed Eng 5, 555-+ (2021). 10.1038/s41551-020-00682-w

10 Xie, W. et al. Prostate Cancer Risk Stratification via Nondestructive 3D Pathology with Deep Learning-Assisted Gland Analysis. Cancer Res 82, 334–345 (2022). 10.1158/0008-5472.CAN-21-2843

11 Phillip, J. M., Han, K. S., Chen, W. C., Wirtz, D. & Wu, P. H. A robust unsupervised machine-learning method to quantify the morphological heterogeneity of cells and nuclei. Nat Protoc 16, 754–774 (2021). 10.1038/s41596-020-00432-x

12. Chen, L. C., Papandreou, G., Kokkinos, I., Murphy, K. & Yuille, A. L. in IEEE Transactions on Pattern Analysis and Machine Intelligence Vol. 40 834–848 (IEEE Computer Society, 2018).

13 Liu, Z. et al. Swin Transformer: Hierarchical Vision Transformer using Shifted Windows. 2021 Ieee/Cvf International Conference on Computer Vision (Iccv 2021), 9992–10002 (2021). 10.1109/Iccv48922.2021.00986

14 Strudel, R., Garcia, R., Laptev, I. & Schmid, C. Segmenter: Transformer for Semantic Segmentation. 2021 Ieee/Cvf International Conference on Computer Vision (Iccv 2021), 7242–7252 (2021). 10.1109/Iccv48922.2021.00717

15 Belevich, I. & Jokitalo, E. DeepMIB: User-friendly and open-source software for training of deep learning network for biological image segmentation. PLoS Comput Biol 17, e1008374 (2021). 10.1371/journal.pcbi.1008374

16 Gomez-de-Mariscal, E. et al. DeepImageJ: A user-friendly environment to run deep learning models in ImageJ. Nat Methods 18, 1192–1195 (2021). 10.1038/s41592-021-01262-9

17 Lutnick, B. et al. A user-friendly tool for cloud-based whole slide image segmentation with examples from renal histopathology. Commun Med (Lond) 2, 105 (2022). 10.1038/s43856-022-00138-z

18 Ghahremani, P., Marino, J., Dodds, R. & Nadeem, S. DeepLIIF: An Online Platform for Quantification of Clinical Pathology Slides. 2022 Ieee/Cvf Conference on Computer Vision and Pattern Recognition (Cvpr 2022), 21367–21373 (2022). 10.1109/Cvpr52688.2022.02071

19 Muller, A. et al. Modular segmentation, spatial analysis and visualization of volume electron microscopy datasets. Nat Protoc 19, 1436–1466 (2024). 10.1038/s41596-024-00957-5

20 Kiemen, A. L. et al. CODA: quantitative 3D reconstruction of large tissues at cellular resolution. Nat Methods 19, 1490–1499 (2022). 10.1038/s41592-022-01650-9

21 Braxton, A. M. et al. 3D genomic mapping reveals multifocality of human pancreatic precancers. Nature (2024). 10.1038/s41586-024-07359-3

22 Kiemen, A. L. et al. Power-law growth models explain incidences and sizes of pancreatic cancer precursor lesions and confirm spatial genomic findings. Science Advances (2024). 10.1101/2023.12.01.569633

23 Dequiedt, L. et al. Three-Dimensional Reconstruction of Fetal Rhesus Macaque Kidneys at Single-Cell Resolution. Laboratory Investigation 103, S1433–S1434 (2023). 10.1101/2023.12.07.570622

24. Forjaz, A. V., E. Matos-Romero, V.; Joshi, S.; Fujikara, K.; Braxton, A.M.; Jiang, A.; Cornish, T.; Hong, S.M.; Hruban, R.H.; Wood, L.; Wu, P.H.; Kiemen, A.; Wirtz, D. Three-dimensional assessments are necessary to determine the true spatial tissue composition of diseased tissues. Biorxiv (2023).

25 Kiemen, A. L. et al. High-Resolution 3D Printing of Pancreatic Ductal Microanatomy Enabled by Serial Histology. Adv Mater Technol-Us 9 (2024). 10.1002/admt.202301837

26 Kiemen, A. L. D., A. I.; Braxton, A.M.; He, J.; Laheru, D.; Fishman, E.K.; Chames, P.; Almagro-Perez, C.; Wu, P.W.; Wirtz, D.; Wood, L. D.; Hruban, R. H. Tissue clearing and 3D reconstruction of digitized, serially sectioned slides provide novel insights into pancreatic cancer. Med (2023). 10.1016/j.medj.2022.11.009

27 Kiemen, A. L. D., L.; Shen, Y; Zhu, Y.; Matos-Romero, V.; Forjaz, A.; Campbell, K.; Dhana, W.; Cornish, T.; Braxton, A.; Wu, P.; Fishman, E.; Wood, L.; Wirtz, D.; Hruban, R. PanIN or IPMN? Redefining lesion size in three dimensions. American Journal of Surgical Pathology (2024). 10.1097/PAS.0000000000002245

28 Johnson, J. A. I. et al. Digitize your Biology! Modeling multicellular systems through interpretable cell behavior. bioRxiv (2023). 10.1101/2023.09.17.557982

29 Joshi, S. et al. Generative interpolation and restoration of images using deep learning for improved 3D tissue mapping. bioRxiv (2024). 10.1101/2024.03.07.583909

30 Sidiropoulos, D. N. et al. Machine learning integrating spatial omics uncovers humoral immunity patterns in intratumoral tertiary lymphoid structures in pancreatic cancer pathologic responders. Cancer Research 84 (2024). 10.1158/1538-7445.Am2024-1159

31 Deshpande, A. L., M.; Sidiripoulos, D. N.; Zhangm S.; Yuanm L; Bell, A.; Zhu, Q. Jin Ho, W.; Santa-Maria, C.; Gilkes, D.; Williams, S. R.; Uytingco, C.R.; Chew, J.; Hartnett, A.; Bent, Z.W.; Favorov, A. V.; Popel, A.S.; Yarchoan, M.; Kiemen, A.; Wu, P.H.; Fujikura, K.; Wirtz, D.; Wood, L.; Zheng, L.; Jaffee, E. M.; Anders, R.; Danilova, L.; Stein-O’Brien, G.; Kagohara, L.T.; Fertig, E. Uncovering the spatial landscape of molecular interactions within the tumor microenvironment through latent spaces. Cell Systems (2023). 10.1016/j.cels.2023.03.004

32 Bell, A. M. J.T.;, Kiemen, A. L. F., K.; Fedor, H.; ambichler, B.; Deshpande, A.; Wu, P.; Sidiropoulos, D.; Erbe, R.; Stern, J.; Chan, R.; Williams, S.; Chell, J.M.; Zimmerman, J.W.; Wirtz, D.; Jaffee, E.M.; & Wood, L. D. F., E.J.; Kagohara, L.T.;. PanIN and CAF Transitions in Pancreatic Carcinogenesis Revealed with Spatial Data Integration. Cell Systems (2024). 10.1016/j.cels.2024.07.001

33 Sneider, A. K., A.; Kim, JH; Wu, PH; Habibi, M; White, M.; Phillip, J.M.; Gu, L.; Wirtz, D. Deep learning identification of stiffness markers in breast cancer. Biomaterials 285 (2022). 10.1016/J.BIOMATERIALS.2022.121540

34 Groot. A. E. d. M., Kayla V.; Krueger, Timothy E. G.; Kiemen, Ashley L.; Nagy, Natalia H.; & Brame, A. T., Vicente E.; Zhang, Zhongyuan; Trabzonlu, Levent; Brennen, W. Nathaniel; Wirtz, Denis; Marzo, Angelo M. De; Amend, Sarah R.; Pienta, Kenneth J. Characterization of tumor-associated macrophages in prostate cancer transgenic mouse models. The Prostate 81, 629–647 (2021). 10.1002/PROS.24139

35 Kiemen, A. L. et al. Three-dimensional immune atlas of pancreatic cancer precursor lesions reveals large inter- and intra-lesion heterogeneity. Cancer Research 84 (2024). 10.1158/1538-7445.Am2024-1206

36 Kiemen, A. L. et al. Intraparenchymal metastases as a cause for local recurrence of pancreatic cancer. Histopathology (2022). 10.1111/his.14839

37 Crawford, A. J. et al. Precision-engineered biomimetics: the human fallopian tube. Science Advances (2024). 10.1101/2023.06.06.543923

38 Lee, M. H. et al. Multi-compartment tumor organoids. Mater Today 61, 104–116 (2022). 10.1016/j.mattod.2022.07.006

39 Xue, Y. et al. Mechanical tension mobilizes Lgr6+ epidermal stem cells to drive skin growth. Science Advances 8, 8698 (2022). 10.1126/SCIADV.ABL8698/SUPPL_FILE/SCIADV.ABL8698_MOVIES_S1_TO_S4.ZIP

40 Yang, H. et al. Engineered bispecific antibodies targeting the interleukin-6 and -8 receptors potently inhibit cancer cell migration and tumor metastasis. Molecular Therapy (2022). 10.1016/J.YMTHE.2022.07.008

41 O’Brien, J. et al. Skin keratinocyte-derived SIRT1 and BDNF modulate mechanical allodynia in mouse models of diabetic neuropathy. Brain (2024). 10.1093/brain/awae100

42 Forjaz, A. K., D.; Shen, Yu; Bea, H.; Tsapatsis, M.; Ping, J.; Queiroga, V; San, K.; Joshi, S.; Grubel, C.; Beery, M.L.; Kusmartseva, I.; Atkinson, M.; Kiemen, A.L.; Wirtz, D. Integration of nuclear morphology and 3D imaging to profile cellular neighborhoods. biorxiv (2025). 10.1101/2025.03.31.646356

43 Montezuma, D. et al. Annotation Practices in Computational Pathology: A European Society of Digital and Integrative Pathology (ESDIP) Survey Study. Lab Invest 105, 102203 (2024). 10.1016/j.labinv.2024.102203

44 Liao, Y. H., Kar, A. & Fidler, S. Towards Good Practices for Efficiently Annotating Large-Scale Image Classification Datasets. Proc Cvpr Ieee, 4348-4357 (2021). 10.1109/Cvpr46437.2021.00433

45 Kirillov, A. et al. Segment Anything. Ieee I Conf Comp Vis, 3992-4003 (2023). 10.1109/Iccv51070.2023.00371

46 Moor, M. et al. Foundation models for generalist medical artificial intelligence. Nature 616, 259–265 (2023). 10.1038/s41586-023-05881-4

47 Zhang, K. et al. A generalist vision-language foundation model for diverse biomedical tasks. Nat Med 30, 3129–3141 (2024). 10.1038/s41591-024-03185-2

48 Chen, R. J. et al. Scaling Vision Transformers to Gigapixel Images via Hierarchical Self-Supervised Learning. 2022 Ieee/Cvf Conference on Computer Vision and Pattern Recognition (Cvpr 2022), 16123–16134 (2022). 10.1109/Cvpr52688.2022.01567

49 Goode, A., Gilbert, B., Harkes, J., Jukic, D. & Satyanarayanan, M. in Journal of Pathology Informatics Vol. 4 27 (Wolters Kluwer -- Medknow Publications, 2013).

50 He, K. M., Zhang, X. Y., Ren, S. Q. & Sun, J. Deep Residual Learning for Image Recognition. 2016 Ieee Conference on Computer Vision and Pattern Recognition (Cvpr), 770–778 (2016). 10.1109/Cvpr.2016.90

51 Zhang, Y. et al. Myeloid cells are required for PD-1/PD-L1 checkpoint activation and the establishment of an immunosuppressive environment in pancreatic cancer. Gut 66, 124–136 (2017). 10.1136/gutjnl-2016-312078

52 Li, H. Y. et al. A comprehensive benchmarking with practical guidelines for cellular deconvolution of spatial transcriptomics. Nature Communications 14 (2023). ARTN 1548 10.1038/s41467-023-37168-7

53 Dong, R. & Yuan, G. C. SpatialDWLS: accurate deconvolution of spatial transcriptomic data. Genome Biology 22 (2021). ARTN 145 10.1186/s13059-021-02362-7

54 Zhou, Z. X., Zhong, Y. S., Zhang, Z. M. & Ren, X. W. Spatial transcriptomics deconvolution at single-cell resolution using Redeconve. Nature Communications 14 (2023). ARTN 7930 10.1038/s41467-023-43600-9

55 Olson, S. R. et al. Thrombosis and Bleeding in Extracorporeal Membrane Oxygenation (ECMO) Without Anticoagulation: A Systematic Review. ASAIO J 67, 290–296 (2021). 10.1097/MAT.0000000000001230

56 Wang, J. S. H. et al. Multimodality Quantification of Thrombus Deposition in Extracorporeal Membrane Oxygenation (ECMO); Correlating Oxygenator Computed Tomography Imaging, Electron Microscopy and Histology to Clinical Outcomes. Blood 142 (2023). 10.1182/blood-2023-191092

57 Si Han Wang, A. A. R., Caleb H Moon, Ari Lauthner, Helen H Vu, Sandra Rugonyi, Anna J Hansen, Heather M Mayes, Bishoy Zakhary, David Zonies, Ran Ran, Akram Khan, Denis Wirtz, Ashley L Kiemen, Owen McCarty, Joseph J. Shatzel. Development of a method for visualizing and quantifying thrombus formation in extracorporeal membrane oxygenators. under review (2025).

