## supplementary file 1 for "CODAvision: best practices and a user-friendly interface for rapid, customizable segmentation of medical images"

*"Annotating at times can be considered an art,*

*nor everything is black, nor white,*

*but whatever you choose to be grey,*

*be sure it stays the same across the whole slide."*

### Annotation Guide

The provided annotation guide presents an overview of best annotation practices that should be followed when generating training datasets for segmentation models using **Aperio Image Scope**. This manual is crafted within the framework of the CODA workflow, a deep learning-based software for microanatomical tissue labelling. CODA owes its high accuracy to multiple factors, but rigorous and reproducible annotation standards are the perhaps the biggest contributor to robust AI models.

#### Section 1 - What is annotating?

Histological annotation involves outlining specific structures in digitized histology slides. The primary objective is to generate a labeled dataset that serves as "ground truth" to train and evaluate a deep learning model, enabling it to learn and discern patterns associated with different tissue classes.

#### Section 2 – Annotating with Aperio ImageScope

Aperio ImageScope, a freeware developed by Leica Biosystems, enables users to visualize and interact with large histological images generated by a microscope slide scanner from glass tissue slides. Note, as of today, ImageScope is only available for Windows.

Within this guide, we describe the process of annotating utilizing Aperio ImageScope [v12.4.3.5008]. ImageScope opens '**.ndpi**', '**.svs**', '**.scn**', '**.jpg**', and '**.ome.tif**' image files and generates '**.xml**' files for storing the corresponding annotations. The '.xml' files will be generated automatically when annotating an image in ImageScope. See **Fig 1**, for an example of an annotated histological image.

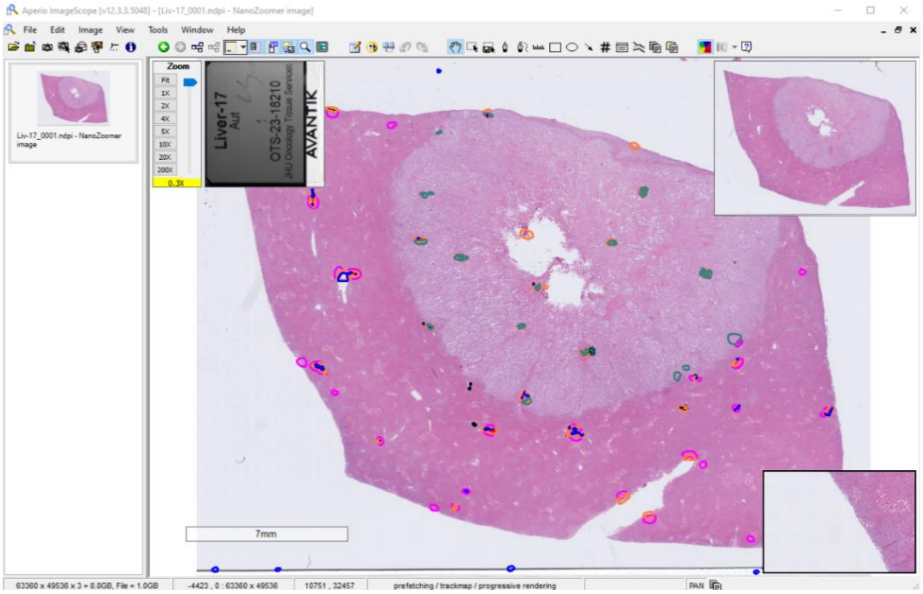


*Figure 1: Example of an annotated human liver whole slide image in ImageScope*

##### 2.1 ImageScope Settings

We recommend adjusting the maximum magnification level allowed within the app to enable enhanced zoom functionality for highly precise annotations. To implement this, navigate to Tools > Options > General Tab > Maximum magnification and modify the value to 1000%. Refer to **Fig 2**.

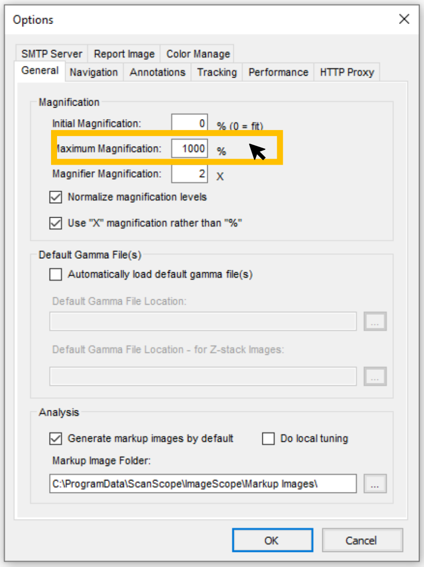


*Figure 2: Preset magnification to 1000%*

We also recommend configuring automatic annotation saving to prevent data loss upon closing the app. To enable this feature, navigate to **Tools > Options > Annotation Tab > Annotation Settings**, and check the box for '**Automatically save annotation changes**,' as illustrated in **Fig 3**. We still recommend saving your annotations periodically as an extra precaution.

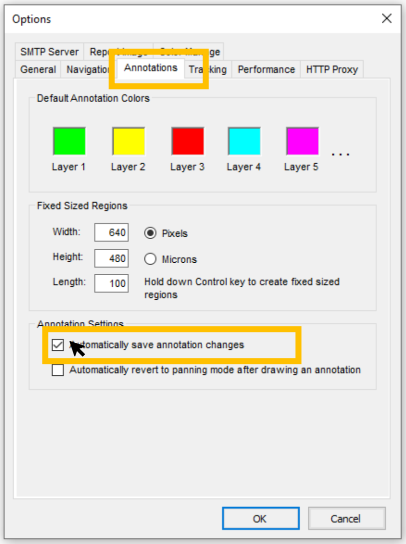


*Figure 3: Configure automatic annotation saving.*

#### Section 3 – How to annotate?

##### 3.1 Building your dataset.

When building the dataset to train your model, start with a **minimum of five whole slide images** for the annotation dataset, although a larger quantity will generally lead to improved results. The selection of these slides should be comprehensive enough to capture the heterogeneity of samples of the project, allowing development of a robust model.

*Note, In the scenario of a sample being a stack of serially sectioned tissue, it is advisable to annotate images from the upper, middle, and lower ends of the stack to create the dataset. See **Fig 4**. Furthermore, attention should be given to the nomenclature of image filenames to ensure correct processing of the images. Avoid using complex symbols such as '-', '*', '.', or blank spaces. An appropriate example of a filename would be 'pancreas_001.ndpi', while an incorrect format would be ''pancreas.tissue*- 001'.

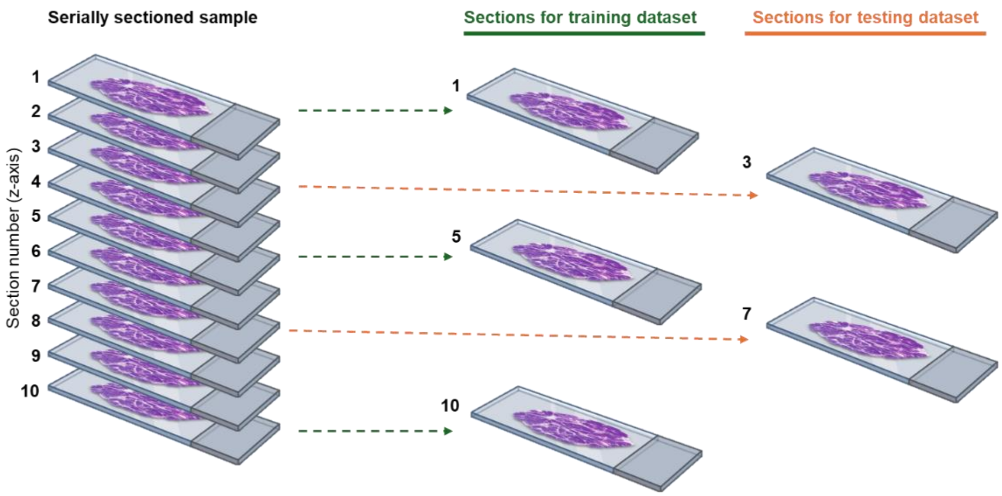


*Figure 4:Training and testing sections for a serially sectioned sample.*

##### 3.2 Training and testing datasets.

To train and assess the performance of any deep learning model, your annotation dataset should be divided into two subsets, one for **training** and another one for **testing**, keeping a ratio around 80% of the annotated images for training and the rest 20% for testing.

The **training dataset** can be defined as the data used to teach the model the different tissue structures it needs to classify automatically.

The **testing dataset** is separate data used to evaluate the model's performance on unseen examples after training. As an example, for the case of having 5 fully annotated whole slide images (WSI), store four WSI in one folder which would be used for training and the remaining one WSI in a separate folder that would be used for testing. Note, the annotated images that you use for testing must have at least one annotation of each of the annotations classes you intend to train your model on. Name the folders as you see fit. Refer to Figure 5 for an illustrated example.

*Note, for the case of training and testing a model on a dataset built with serially sectioned samples the testing subset should comprise sections situated at a reasonable distance within the stack from the images used for training. As an example, for a serially sectioned sample with 100 sections, if sections in positions 1, 50, and 10 are chosen for training, sections in position 25 and 75 would be suitable choices for testing, as they lie between two training sections but are not excessively proximate to either. See **Fig 5**.

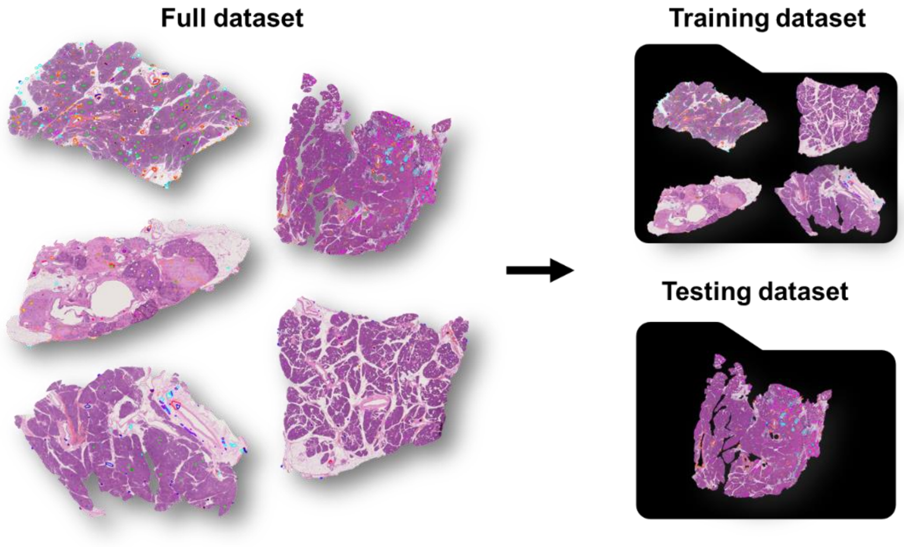


*Figure 5: Training and testing dataset partition.*

##### 3.3 Annotate with Aperio ImageScope

3.3.1 Defining annotation classes.

Selection of annotation classes is closely tied to the research objectives. Each research project circles around a specific question, and it is this question that shapes the level of detail in distinguishing tissue features during annotating. Where the goal is exhaustive labelling of all subtle cell types in an image, a high-performance model will be time consuming to obtain but will be highly detailed. For example, see **Fig 6a** where 17 structures of the fetal kidney were detected in H&E. In contrast, where the biological question is targeted, a very limited set of structures may be defined, and model training will be streamlined. See **Fig 6b**, where the same kidney section was segmented into only two labels: vasculature and non-vasculature.

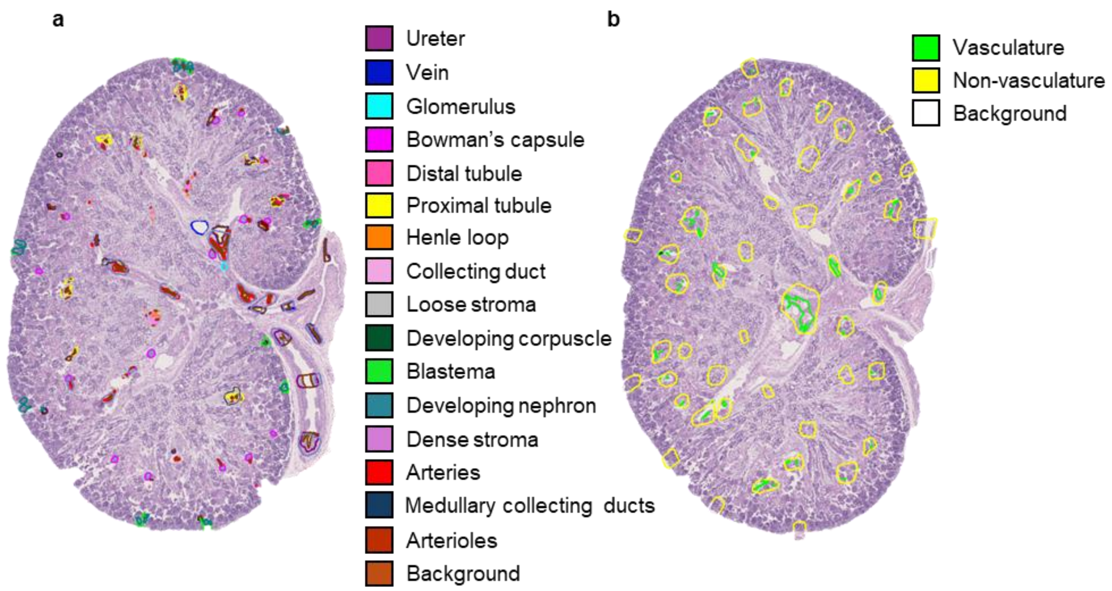


*Figure 6: Annotated pancreas WSI with corresponding labels.*

To avoid misclassification of the tissue structures once the model is trained, all discernible tissue structures should be annotated into an annotation class. Otherwise, the model would get confused with the unannotated structures during the classification step. For example, in the above kidney image, the user should not annotate only 'Glomerulus' and 'Arterioles', as numerous other structures will be unknown to the model. Instead, annotate 'Glomerulus', 'Arterioles', and 'Other kidney structures', with this third class serving as a catch-all for all other structures present in the whole slide images.

##### 3.3.2 Create annotation layers in ImageScope

Once the tissue structures have been defined and the selection of whole slide images to annotate is done, the next step is creating labels for annotations in ImageScope using the 'Annotations - Detailed View' window. To access this window, navigate to **View > Annotations** in the control panel or utilize the shortcut (**Ctrl + N**). This window serves as the interface for generating annotation labels and assigning corresponding colors.

To create a new annotation layer, click the '**+**' icon within the interface (refer to **Fig 7**), which will automatically add an annotation class or layer. ImageScope assigns a default color to the created layer, which can be modified by selecting the color box in the upper section of the window and choosing the preferred color. To alter the name of the annotation class layer, click on the layer name's top and press '**F2**,' then input the desired name. Repeat this procedure to create as many annotation layers and classes as necessary. Information regarding the annotation layers and annotated regions is stored in an '.xml' file with the same name as the annotated image. This file will be created automatically by ImageScope once you start annotating.

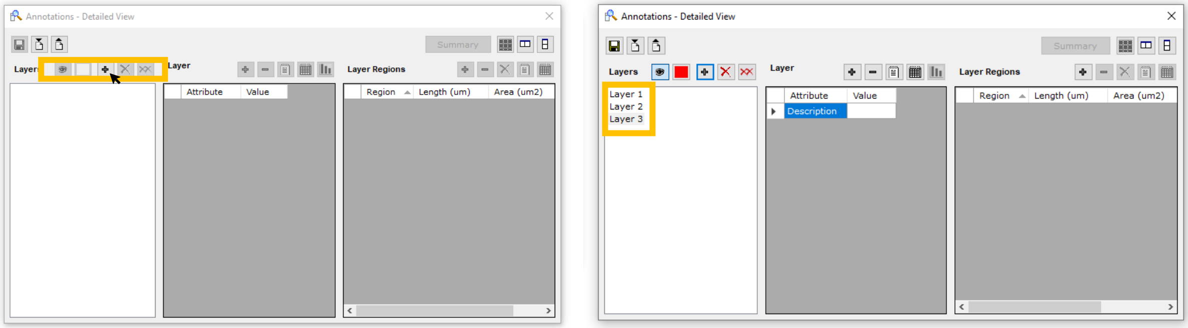


*Figure 7: Operating with layers in ' Annotations – Detailed View' window (ImageScope)*

*Note, select annotation class colors that are easily distinguishable from one another. These colors will serve as default for the classification once your CODA model is trained. However, it is possible to modify these colors later in the protocol if desired (refer to 'CODA Dashboard Guide').

##### 3.3.3 Annotation template

When training a CODA model, the order of annotation layers in Aperio ImageScope holds significance is crucial and must be maintained across all annotated images. To maintain a consistent order of layers across all annotated images within the dataset, create a blank '**.xml**' file following the process explained in Section 3.3.2 for any image in your dataset. Add all annotation layers and save the file.

Next, rename the '**.xml**' file ImageScope created to '**Template.xml**'. Copy and paste this file and rename it to the same name of the image you plan to annotate. This will ensure that when you open the image on Aperio ImageSope, all annotation layers and colors are already predefined. Repeat this process for all the images you plan on annotating.

Alternatively, you can open the '**Annotations - Detailed View**' window in Image Scope and create the layers from scratch for each of the images you annotate, just keep in mind that the order of the layers should be the same for all annotated images, even when one or more annotation layers are not present in the image

##### 3.3.4 Free-form drawing

To annotate, select the '**Free-form drawing**' modality using the '**Pen**' tool in ImageScope. To access this tool, click 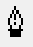 on the toolbar or use the shortcut pressing **F2**.

Next, begin the annotation by clicking and dragging around the area of interest in the main window. Use the pen tool to manually outline the region of interest, closing the drawn shape once completed. If you wish to modify the annotation, click and redraw over the annotated region until achieving the desired level of precision in refining the borders of the annotated area. The annotated region would be added to the annotation class selected in the '**Annotation - Detailed View**' pop up window. See **Fig 8** as reference.

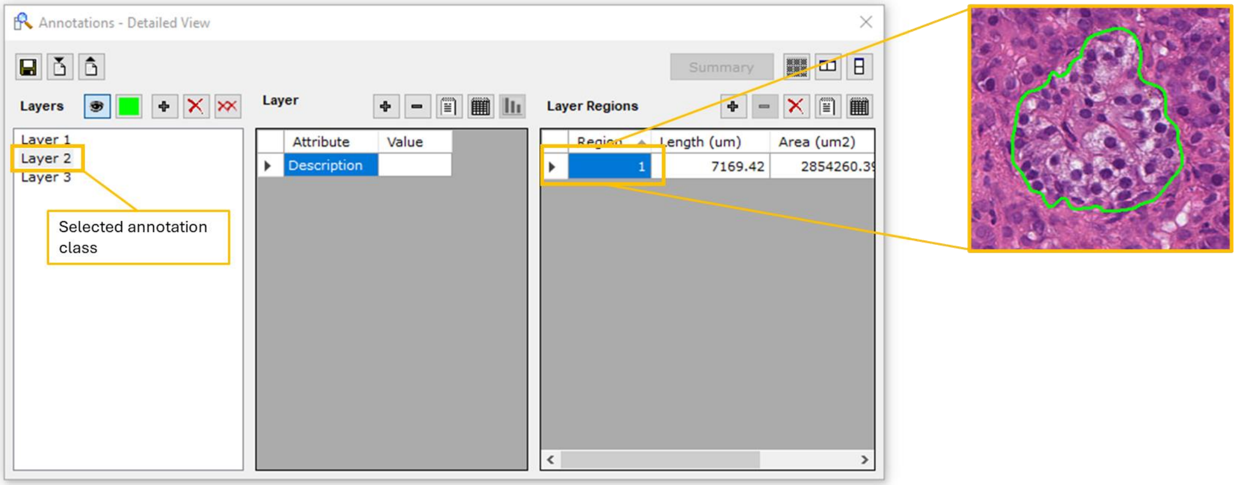


*Figure 8: Annotated region using Pen Tool (Aperio ImageScope)*

#### 3.3.5 Nesting

Our results suggest that segmentation models yield better results when the tissues are identified within their microenvironments. To achieve this, we suggest annotation technique called 'nesting'. Nesting is based in a hierarchical tissue organization, where higher level tissues can be 'nested' inside lower-level ones. Refer to the example in **Fig 9**.

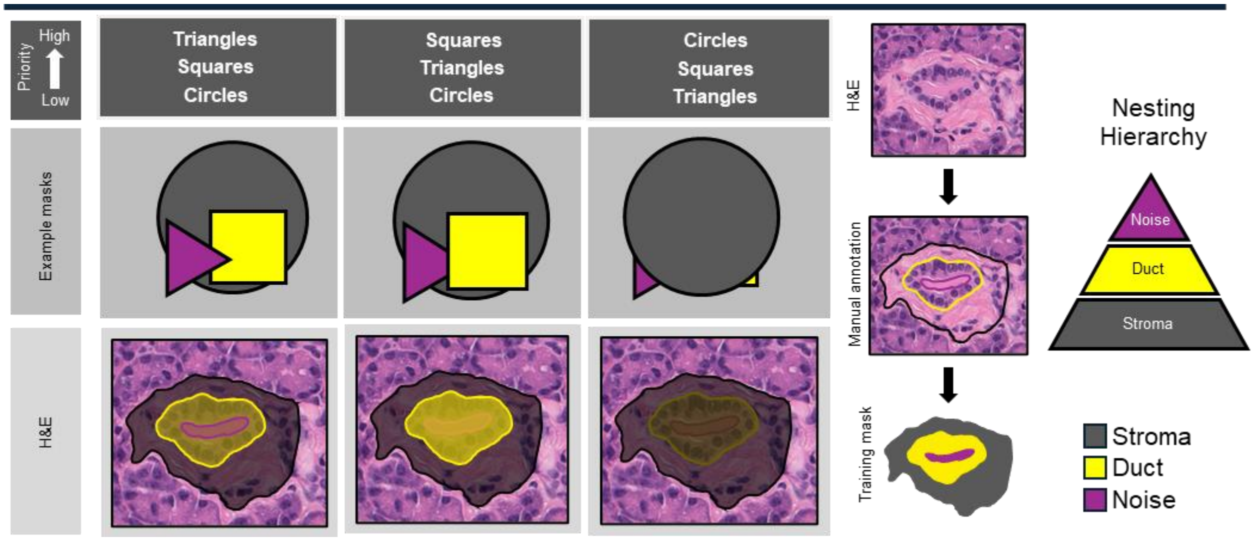


*Figure 9: Annotation hierarchy and quality control for CODA deep learning model training.*

The example provided in **Fig 9** illustrates the arrangement of three hypothetical annotated tissue types: tissue A (triangles), tissue B (squares), and tissue C (circles). It can be observed that tissue A annotations often overlap with annotations from the other two tissues and are generally smaller in size. Similarly, tissue B can be found surrounded by tissue C.

Based on these observations, a nesting hierarchy for this image can be established. Consider the example where tissue A is assigned to the top layer, tissue B to the middle layer, and tissue C to the bottom layer. With this hierarchy, the annotation class for tissue A at the top layer implies that all tissue pixels inside that annotation will be categorized as tissue A. The pixels between the tissue A annotation and the tissue C annotation will be classified as tissue B. Lastly, the pixels within the tissue C annotation but outside both tissue B and A will be categorized as tissue C.

Please note that the hierarchy chosen in cases where annotations overlap is not a parameter that needs to be defined in ImageScope; instead, it will be later specified in the CODA Vision GUI before training the model. Nevertheless, it is essential for the annotator to bear this hierarchy in mind to ensure consistency when dealing with overlapping annotations.

**Fig 10** shows several examples of overlapping annotation for different tissue types.

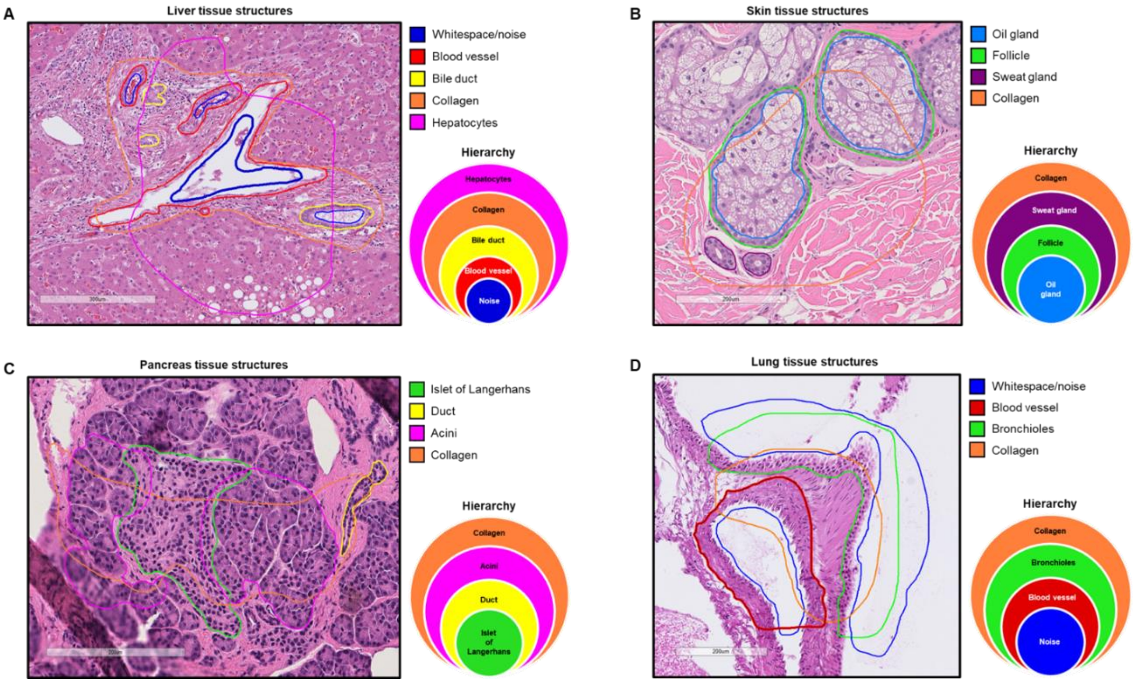


*Figure 10: Nesting examples. A. Overlapping annotations for liver tissue structures. B. Overlapping annotations for skin tissue structures. C. Overlapping annotations for pancreas tissue structures. overlapping annotations for lung tissue structures.*

##### 3. 7 Good Annotation Practices

When creating the annotation dataset, your aim should be to strike a balance between data volume, annotation quality, and the requirements of the target analysis or model. Developing a well-thought-out annotation strategy, aligned with the goals of your study and the characteristics of the annotated objects, will contribute to obtaining reliable and meaningful results. The next concepts comprise various strategies that we suggest following to achieve an optimal dataset. This, in turn, leads to effective training of the deep learning model, saving considerable trial-and-error time.

- **Define Annotation Classes:** Clearly define annotation classes with distinct morphological differences. Well-defined classes enhance the quality of the semantic segmentation by the model.
- **Color Coding and Labeling:** Employ consistent color coding among the different annotated classes. Clearly label annotations with legible text to facilitate identification of structures. See **Fig 11**.

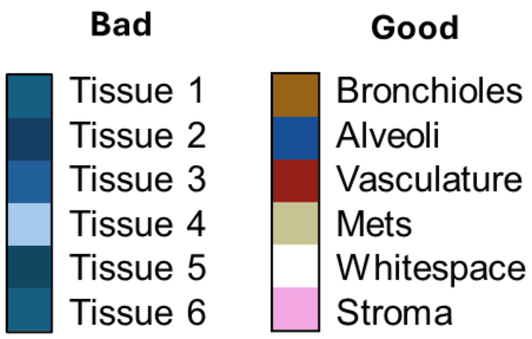


*Figure 11: Optimal labeling examples*

- **Number of Annotations:** The quantity of annotated structures required per class may vary based on your dataset volume, the complexity of structures, specificity of classification, and study goals. To meet the deep learning model's requirements for the annotated dataset used in training, it is advisable to annotate approximately 20-25 structures per defined class in your analysis. This approach builds a substantial dataset foundation, capturing the diversity and variability of the structures under analysis, and contributes to the development of a robust CODA CNN model**.** See **Fig 12**.

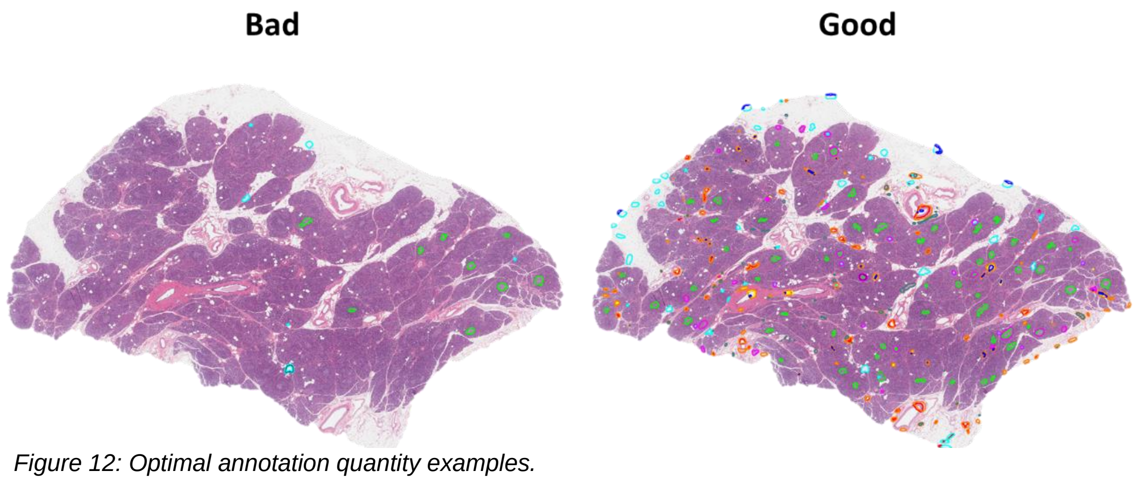


*Figure 12: Optimal annotation quantity examples.*

- **Annotation Heterogeneity:** Building an annotation dataset with a diverse range of morphologies for each annotation class ensures optimal performance of your CODA model on unseen data. In the scenario of applying your model to a dataset with a diverse range of shapes and sizes, a higher number of annotated instances is required to account for this variability. See **Fig 13**.

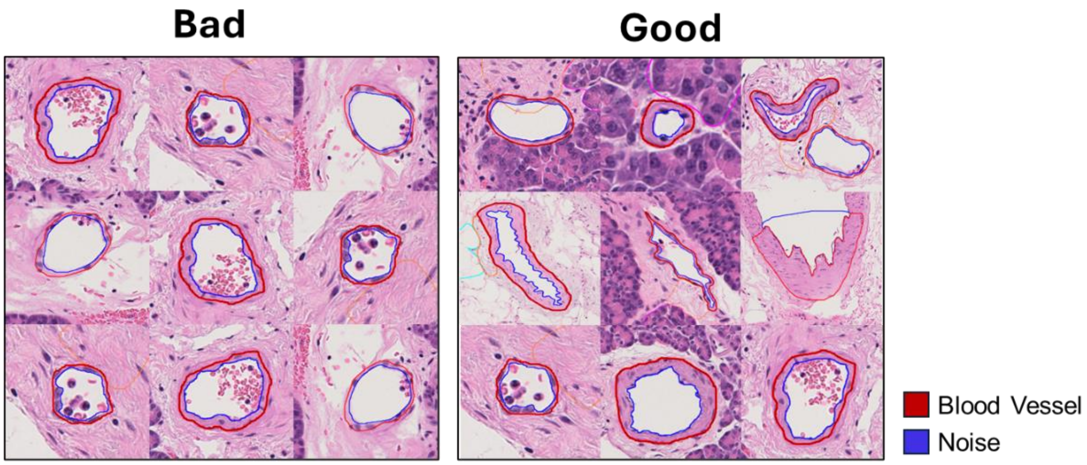


*Figure 13: Optimal annotation heterogeneity examples of a blood vessel. Inadequate annotations on the left are too homogeneous and not property capture all the range of sizes and shapes present in blood vessels. Good annotations from the right offer a diverse array, enhancing heterogeneity.*

- **Noise and Artifacts:** In whole slide images, it's common for tissue to fold over itself during the sectioning process, leading to the appearance of artifacts or noise atop the structures of interest. To enhance your model's performance in handling these defects, incorporate annotations in each of your defined tissue classes, including a few instances with artifacts. This approach enables the model to learn proper classification of the tissue class even in the presence of artifacts or noise. See **Fig 14**. See Section 4 for a more detailed description of noise annotations.

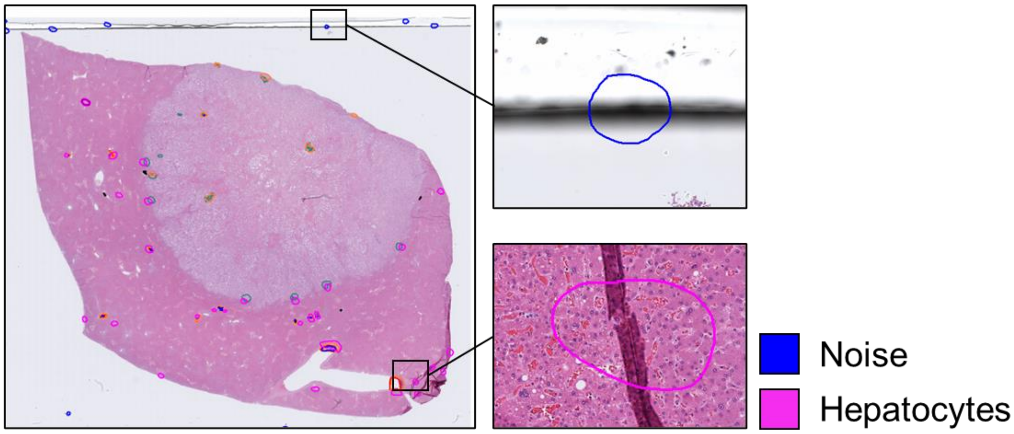


*Figure 14: Noise annotation examples.*

- **Precision and detail:** Be precise in outlining the boundaries of your annotated structures. For CODA, it is recommended to keep an adequate <u>minimum zoom level</u> during annotating clear and precise borders of the annotation, as depicted in **Fig 15**. Note, a model can only be as good as the annotations it is fed with. Adding unprecise annotations to your dataset can only lead to an unprecise classification by your model.

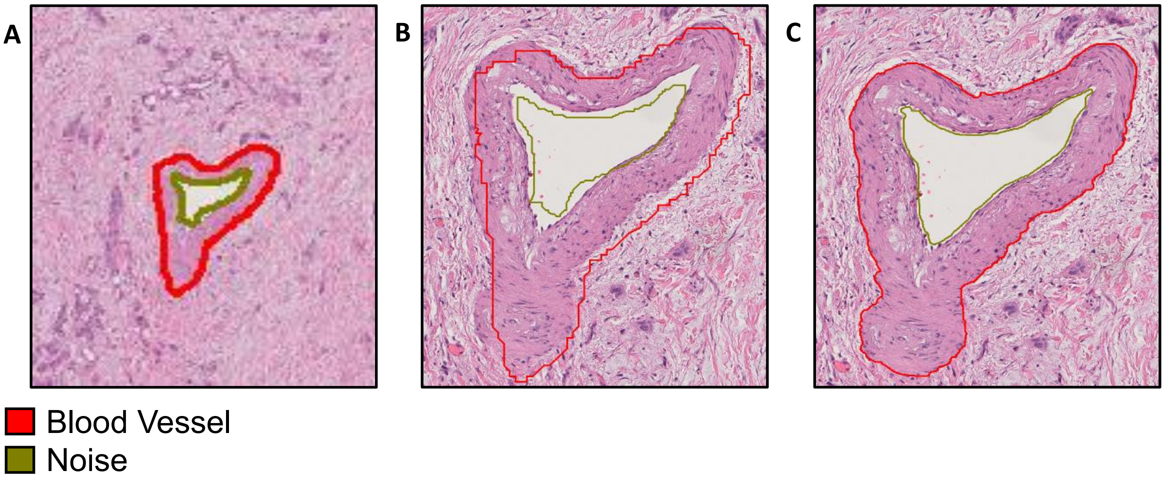


*Figure 15: Comparison of an annotation of a blood vessel made with an inadequate and an optimum zoom level. **A:** Annotation made while Zoomed-out. **B:** Zoom-in of the bad annotation. **C:** Good annotation of the blood vessel*

- **Consistency**: Ensure a uniform annotation style across the dataset by maintaining consistent sizes for your annotations throughout the entire image. Keep a steady threshold for the smallest and largest sizes of the structures you are annotating. See **Fig 16**.

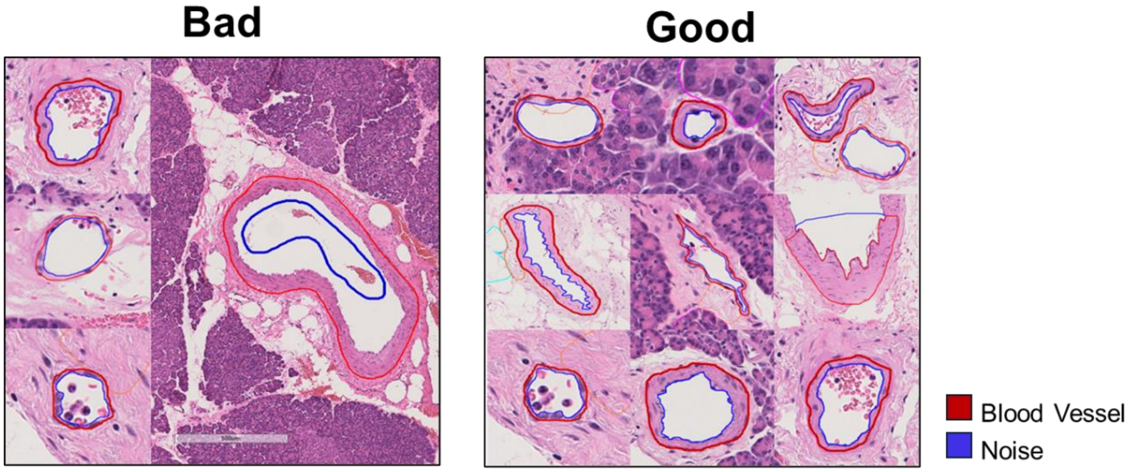


*Figure 16: Comparison of blood vessel annotations using inadequate and optimal annotation styles. Inadequate annotations on the left depict both excessively small and excessively large annotations. Conversely, optimal annotations on the right maintain a consistent average size across annotations while preserving the heterogeneity among the various objects chosen for annotation.*

- **Nesting:** Try to overlap annotations as much as possible. The overlapping of annotations from different classes provides your CODA model with additional context during training, offering insights into the tissue structures surrounding each annotation. This enhances the effectiveness of training and contributes to increased accuracy in classification. See **Fig 17**.

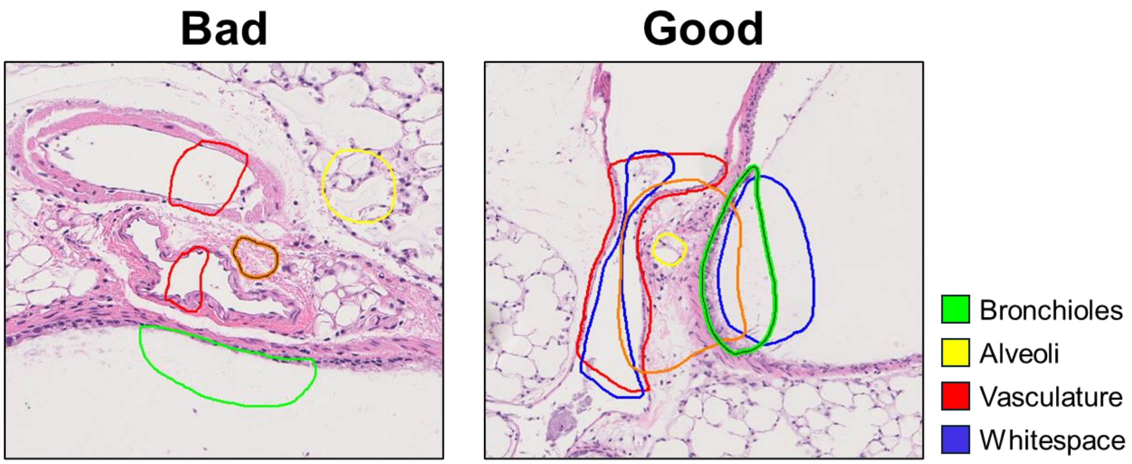


*Figure 17: Comparison of lung tissue annotations with and without nesting. Bad annotations on the left depict 4 different annotation labels annotated separately without nesting. Conversely, good annotations on the right depict the same annotation labels annotated while being nested (overlapped) offering a more comprehensive arrangement of the tissues for the posterior training of the model.*

- **Iterative Approach:** After generating a substantial number of annotated objects in your dataset, initiate model training. Assess the model's performance and increase the number of annotations per class as necessary. Iterate this process until you achieve the desired accuracy and performance levels in your model. **Fig 18** depicts an example of the first classification done by a pilot model build for pancreatic tissue (left) vs the classification of the same tissue slide using a refined model, where additional annotations were incorporated in regions initially misclassified by the preliminary model (right).

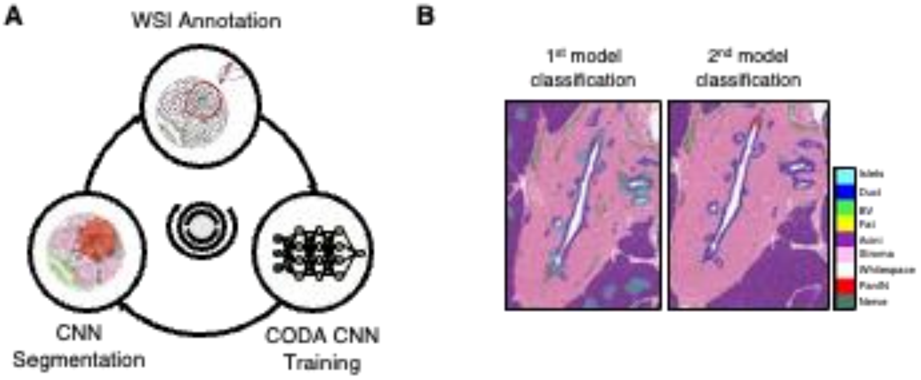


*Figure 18: A. Iterative approach pipeline. B. (Left) Pilot model classification of pancreatic tissue slide. (Right) Second classification after increasing the annotations on misclassified regions.*

#### Section 4 – Noise, Whitespace, and Background recognition

One drawback of tissue sectioning is the potential detachment of cells from their native tissues and the presence of remnant fluids and chemicals in the lumen of ducts. To limit the effect of these artifacts on your deep learning model, introduce a top hierarchy category named "noise/whitespace". The addition of this annotation class aims to train the model to disregard noise objects during classification.

##### 4.1 Lumen Noise

The lumen within blood vessels and ductal structures may contain cells, floating particles, or other forms of noise. When the model classifies this lumen, it may become confused and categorize these objects as the surrounding tissue. To avoid this, annotate the lumen inside these ductal structures as noise. In the illustrated example in **Fig 19**, we can see a pancreatic duct containing noise in the luminal space. This annotation is made to prevent the model from classifying the noise as ductal epithelium.

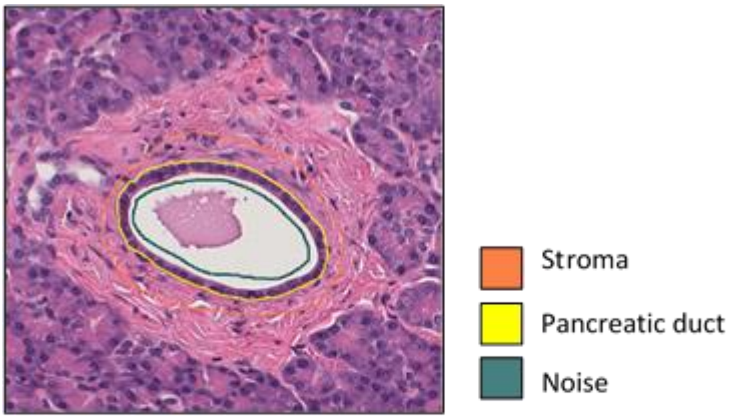


*Figure 19: Pancreatic duct with blood inside annotated as noise.*

##### 4.2 WSI border artifacts

The model might encounter confusion around the slide borders, as the shadow of the coverslip creates a distinctive line in the image. To prevent misinterpretation of artifacts in this space, noise annotations should be added here to ensure proper training when encountering such structures. Include 5 - 10 edge annotations per annotated slide to create a thorough dataset. See **Fig 20**.

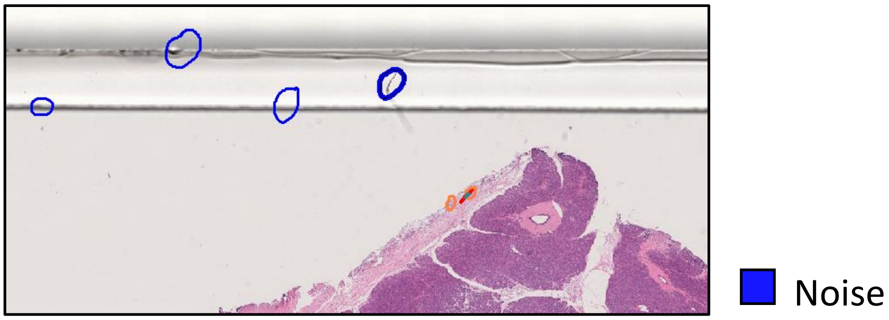


*Figure 20: Noise annotations on artifacts located at the edges of the WSI*

##### 4.3 Edge annotations

To guarantee the accurate differentiation of tissue borders from whitespace during model classification, include annotations at the edges of the tissue. For example, stroma at the edge of tissue is often misclassified as blood vessels, while acinar cells in pancreatic histology at the edge of the tissue are often misclassified as epithelial cells, as these structures are expected to contain an open lumen. We recommend incorporating a minimum of 5-10 tissue edge annotations per annotated slide. See **Fig 21**.

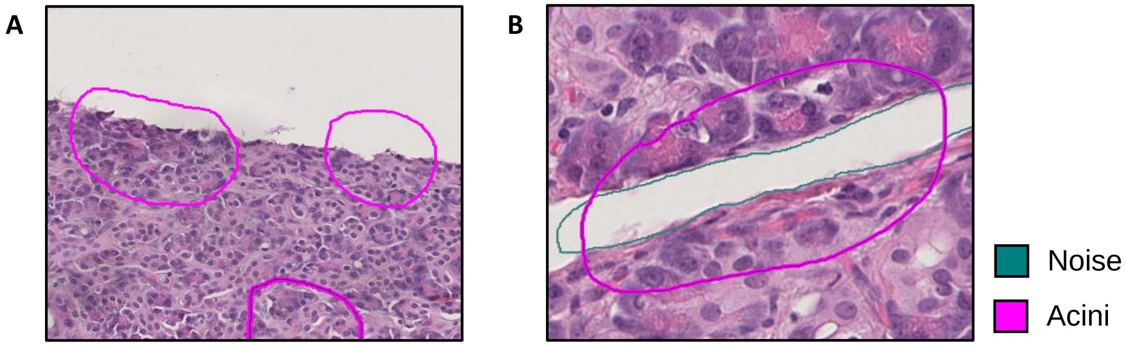


*Figure 21. A: Annotation of acinar cells in the of a slide. B: Annotation of acinar cells at the two sides of a teared slide, overlapping with a noise annotation of the whitespace to avoid misclassification.*

#### Pancreas

| Sub-tissue structure | Description | Annotation | Nesting |
|---|---|---|---|
| Langerhans Islets | Rounded clusters of endocrine cells producing hormones, distinct in staining from surrounding tissue. | 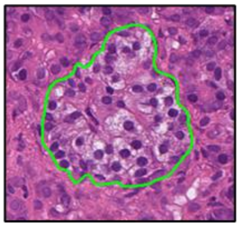 | 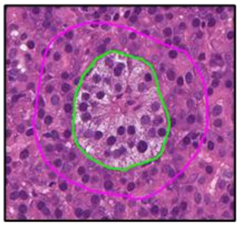 |
| Duct | Tubular structures with a single layer of epithelial cells and a clear lumen. | 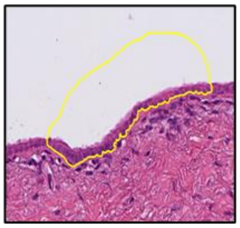 | 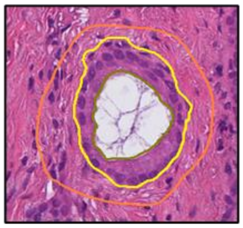 |
| Vasculature | Arteries (thicker) and veins (thinner) appear as circles or ovals with erythrocytes and lumen in center. | 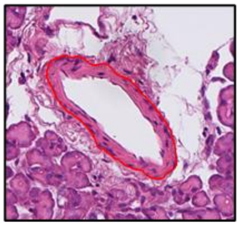 | 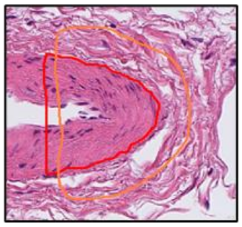 |
| Nerves | Thin, elongated structures with a wavy or branching pattern, often surrounded by connective tissue | 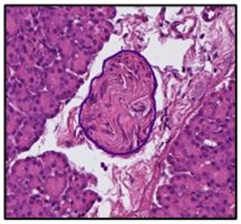 | 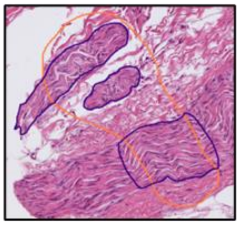 |
| Acini | Grape-like clusters of exocrine cells with a central lumen, darkly stained due to enzyme granules. | 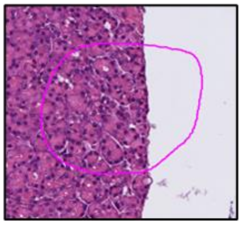 | 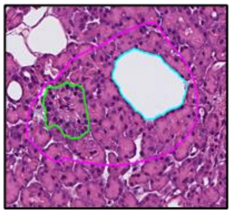 |
| Fat | Adipose tissue with large, round cells that have a central vacuole. | 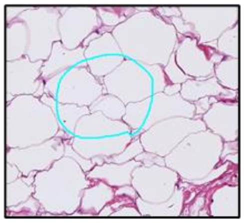 | 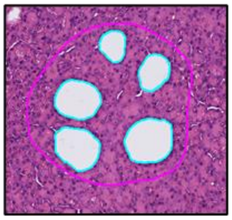 |
| Collagen | Pink or red fibrous tissue forming dense bundles or meshes. | 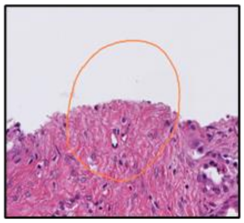 | 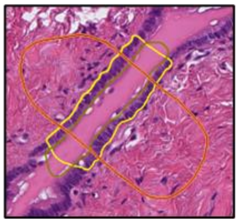 |

#### Pancreas

| Sub-tissue structure | Description | Annotation | Nesting |
|---|---|---|---|
| Whitespace | Areas lacking staining or cellular structures, appearing as empty spaces or noise. | 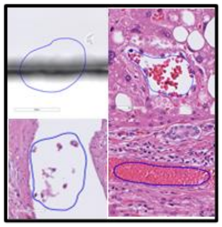 | 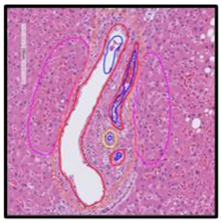 |
| PanIN | Pre-cancerous lesions with abnormal, dysplastic glandular architecture and atypical cells. | 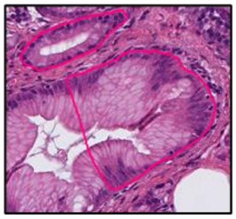 | 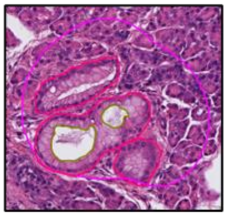 |

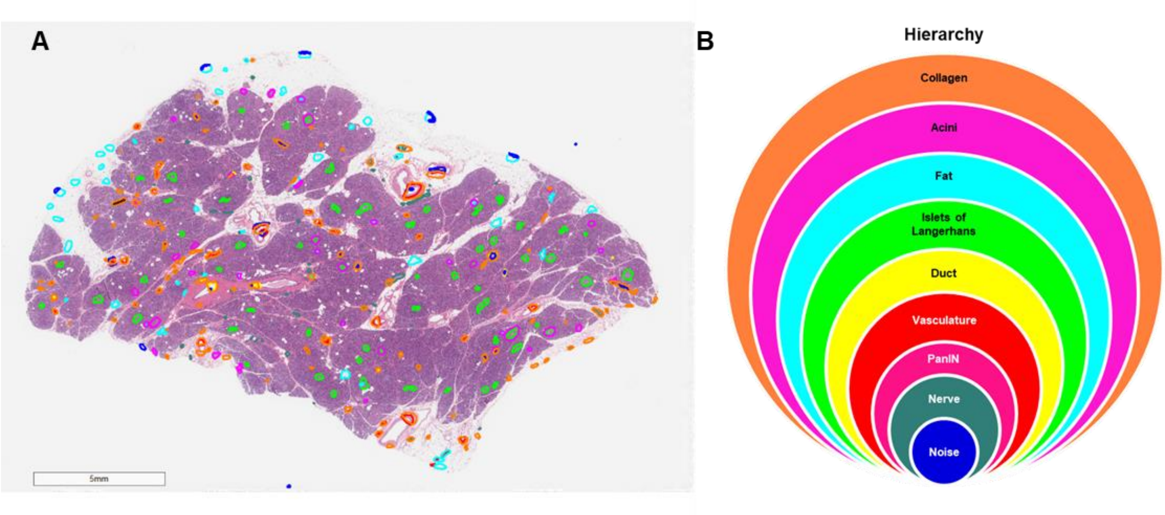


*Figure 22: A. Annotated WSI of pancreatic tissue. B. Nesting hierarchy for the pancreatic tissue structures annotated: 'Noise>Nerve>PanIN>Vasculature>Duct> Islets> Fat> Acini>Collagen'.*

#### Liver

| Sub-tissue structure | Description | Annotation | Nesting |
|---|---|---|---|
| Hepatocytes | Large, polygonal liver cells with central nuclei, organized in cords or plates and separated by blood sinusoids. | 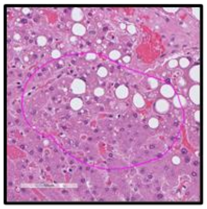 | 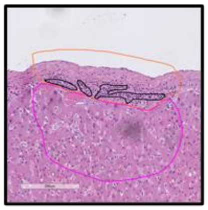 |
| Bile duct | Small, tubular structures with a single layer of cuboidal or columnar cells. | 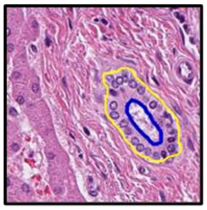 | 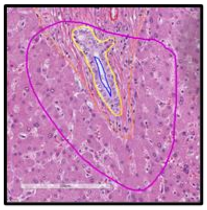 |
| Vasculature | Arteries (thicker) and veins (thinner) appear as circles or ovals with erythrocytes and lumen in center. | 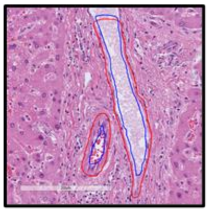 | 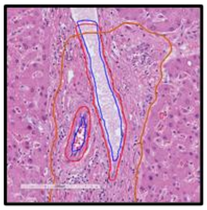 |
| Collagen | Pink or red fibrous tissue forming dense bundles or meshes. | 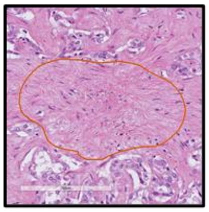 | 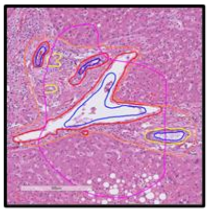 |
| Whitespace | Areas lacking staining or cellular structures, appearing as empty spaces or noise. | 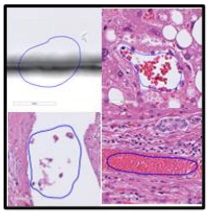 | 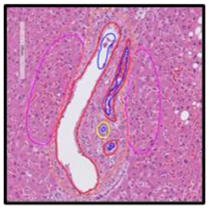 |
| Immune hotspots | Clusters of inflammatory or immune cells, denser than surrounding tissue. | 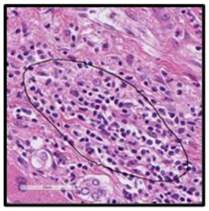 | 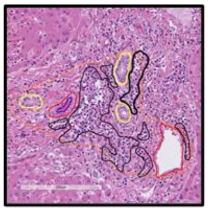 |
| Metastasis | Irregular, invasive glandular structures with atypical cell arrangement and frequent mitosis. | 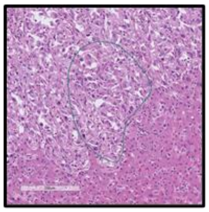 | 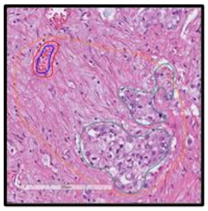 |

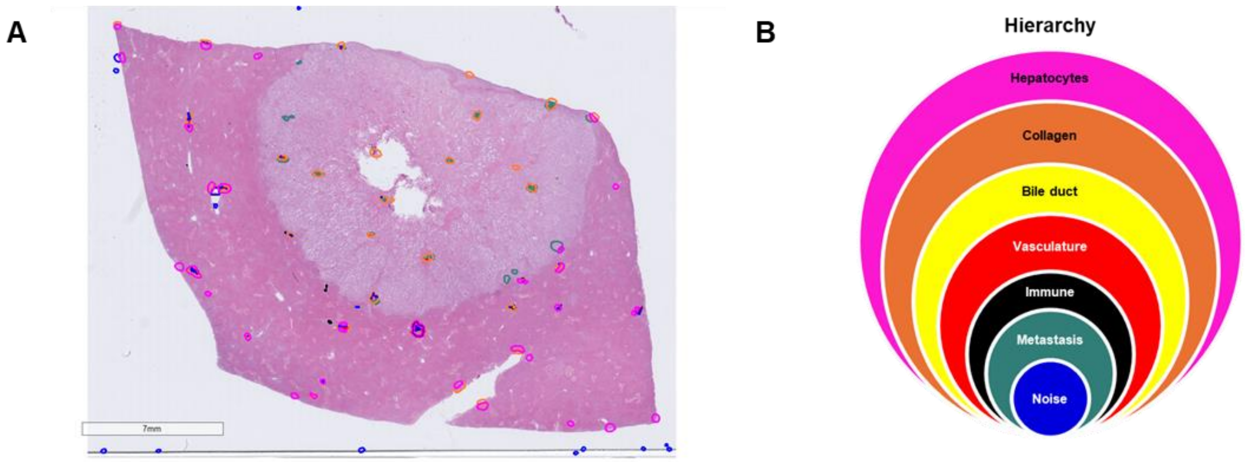


*Figure 23: Figure 7: A. Annotated WSI of liver tissue. B. Nesting hierarchy for the liver tissue structures annotated: 'Noise>Metastasis>Immune>Vasculature>Duct> Hepatocytes> Collagen'.*

#### Skin

| Sub-tissue structure | Description | Annotation | Nesting |
|---|---|---|---|
| Epidermis | Top layer of skin, consisting of several layers of tightly packed epithelial cells | 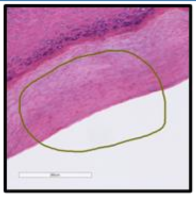 | 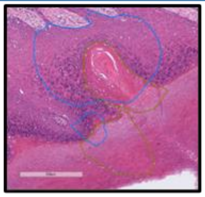 |
| Stratum Corneum | Outermost layer of skin, composed of several layers of tightly packed epithelial cells | 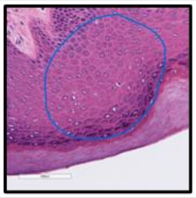 | 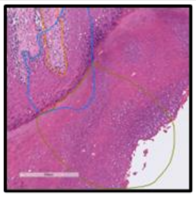 |
| Eccrine glands | Tubular structures with wider, more rounded cells and a lighter staining appearance involved in sweat production. | 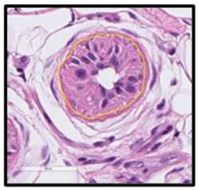 | 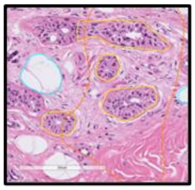 |
| Eccrine ducts | Thin ducts connecting eccrine glands to the skin surface. They are more basophilic (darker staining) in HE. | 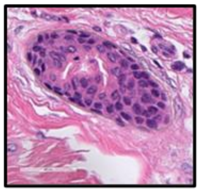 | 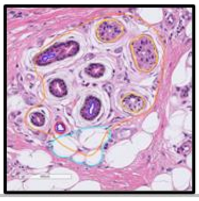 |
| Vasculature | Arteries (thicker) and veins (thinner) appear as circles or ovals with erythrocytes and lumen in center. | 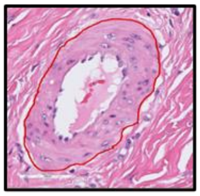 | 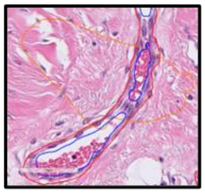 |
| Nerve | Thin, elongated structures with a wavy or branching pattern, often surrounded by connective tissue | 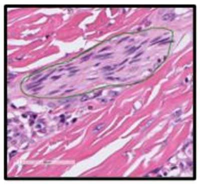 | 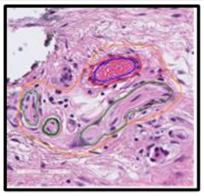 |
| Collagen | Pink or red fibrous tissue forming dense bundles or meshes. | 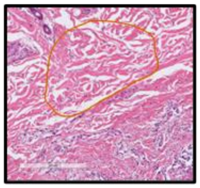 | 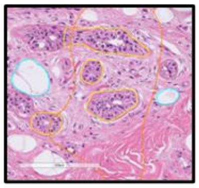 |
| Fat | Adipose tissue with large, round cells that have a central vacuole, commonly found in subcutaneous layers. | 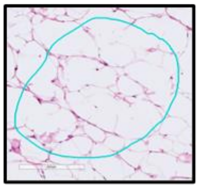 | 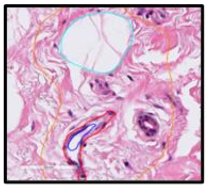 |

| Skin | | | |
|---|---|---|---|
| Sub-tissue structure | Description | Annotation | Nesting |
| Whitespace | Areas lacking staining or cellular structures, appearing as empty spaces or noise. | 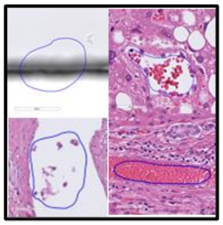 | 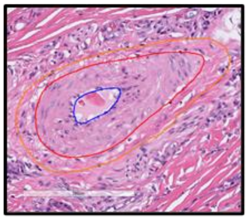 |
| Immune hotspots | Clusters of inflammatory or immune cells, denser than surrounding tissue. | 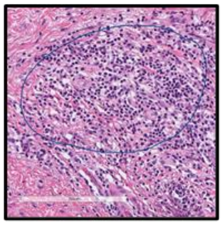 | |

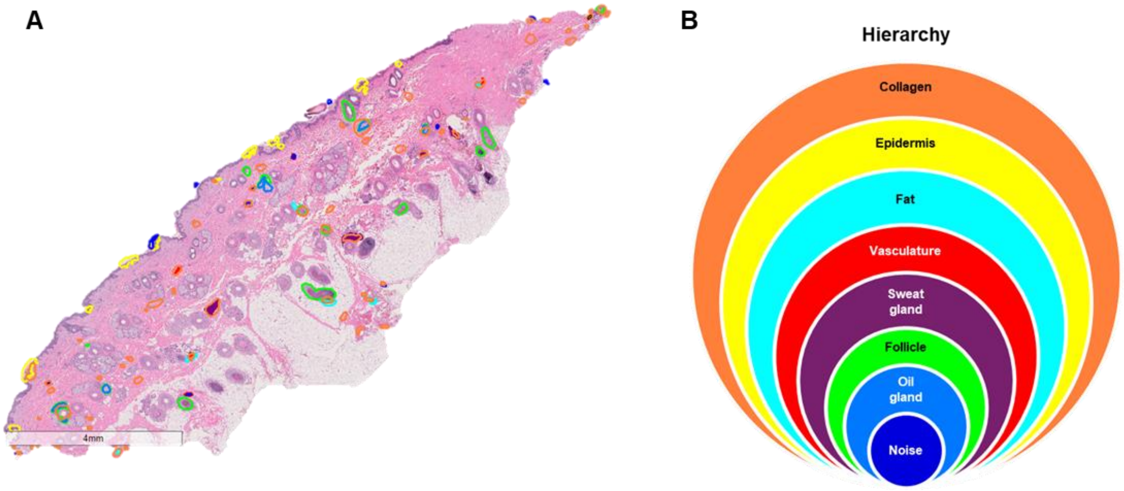


*Figure 24: Annotated WSI of skin tissue. Nesting hierarchy for the skin tissue structures annotated: 'Noise>Oil gland>Follicle>Sweat gland>Vasculature>Fat>Epidermis>Collagen'.24*

#### Lung

| Sub-tissue structure | Description | Annotation | Nesting |
| --- | --- | --- | --- |
| Bronchioles | Thin-walled airways with a smooth muscle layer, lacking cartilage and lined by a simple columnar or cuboidal epithelium. | 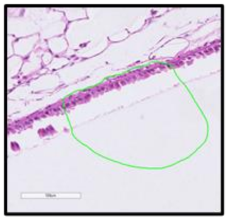 | 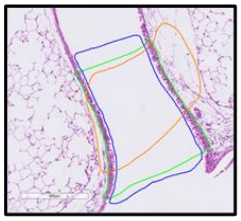 |
| Alveoli | Tiny, sac-like structures with thin walls where gas exchange occurs, lined by a simple squamous epithelium. | 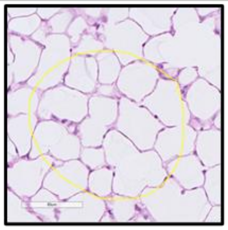 | 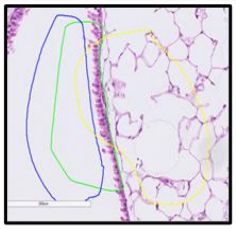 |
| Vasculature | Arteries (thicker) and veins (thinner) appear as circles or ovals with erythrocytes and lumen in center. | 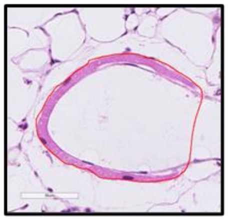 | 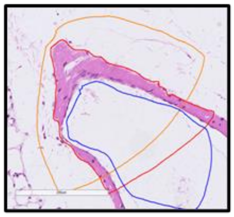 |
| Collagen | Pink or red fibrous tissue forming dense bundles or meshes. | 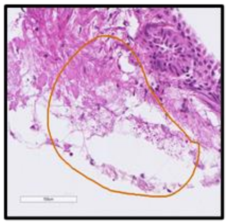 | 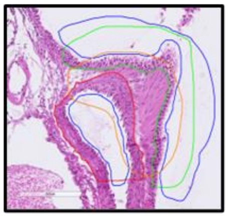 |
| Whitespace | Areas lacking staining or cellular structures, appearing as empty spaces or noise. | 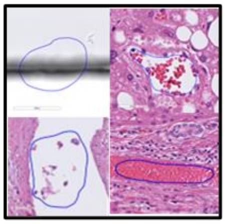 | 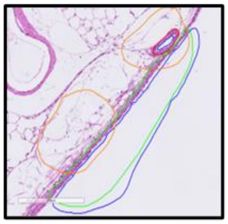 |
| Metastasis | Irregular, invasive glandular structures with atypical cell arrangement and frequent mitosis. | 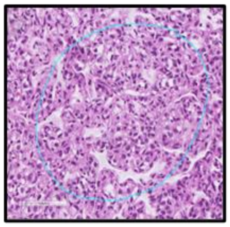 | 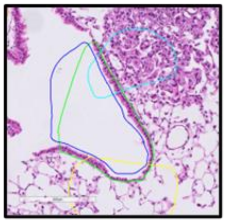 |

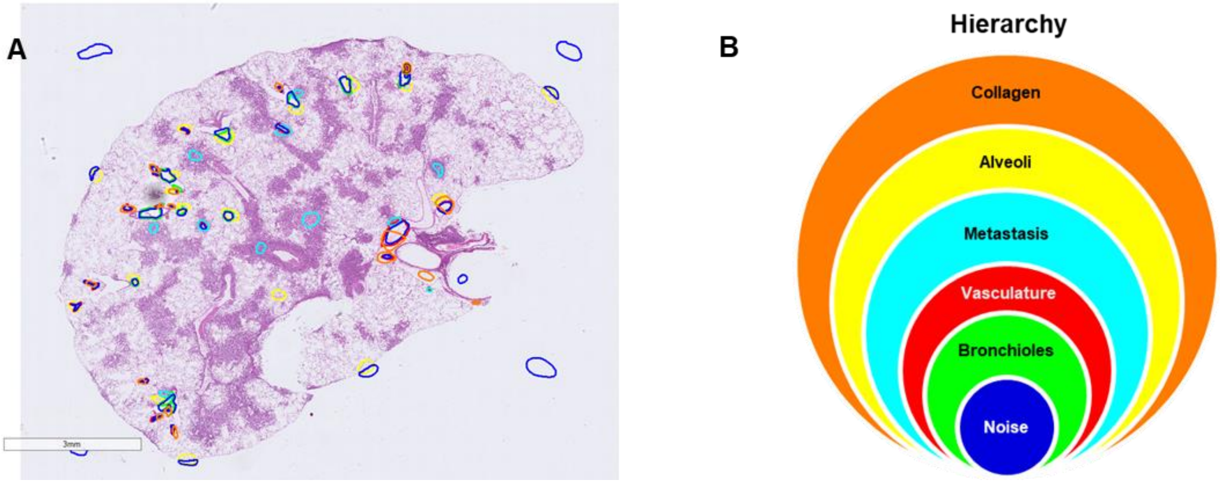


*Figure 25: Annotated WSI of skin tissue. Nesting hierarchy for the skin tissue structures annotated: 'Noise>Bronchioles>Vasculature>Metastasis>Alveoli>Collagen'.*

#### Torso CT

| Sub-tissue structure | Description | Annotation | Nesting |
|---|---|---|---|
| Bone | Dense, bright white structures. The ribs surround the lungs and the spine can be observed at the bottom of the image. | 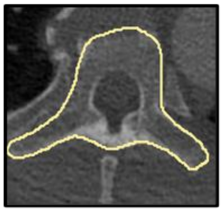 | 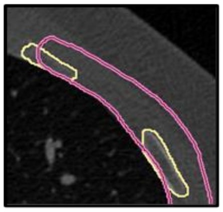 |
| Lung | Air-filled structures, very dark or black, containing smaller light gray areas (blood vessels). | 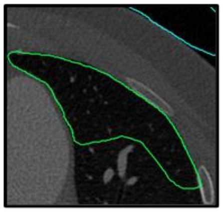 | 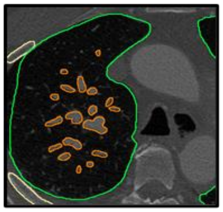 |
| Lung vessels | Small vessels within the lung. They appear as small, light gray structures against the black background of the lung. | 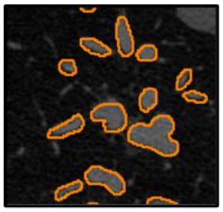 | 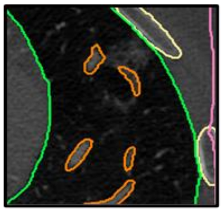 |
| Heart vessels | Major blood vessels like the aorta and pulmonary arteries. Appear as light gray tubular or round structures. | 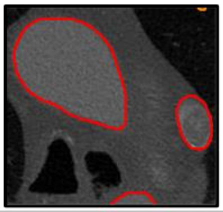 | 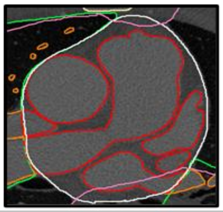 |
| Heart | A dense dark gray structure with lighter areas inside it (blood vessels). It is located between the lungs. | 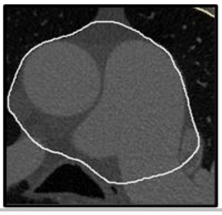 | 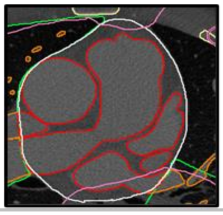 |
| Other Tissues | Tissue areas that do not belong to any of the other classes. | 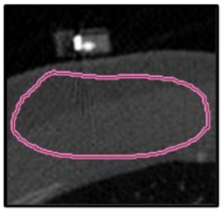 | 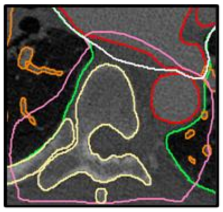 |
| Non-tissue | Areas lacking staining or cellular structures, appearing as empty spaces or noise. | 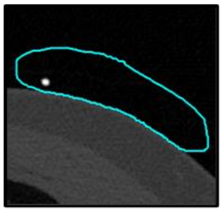 | |

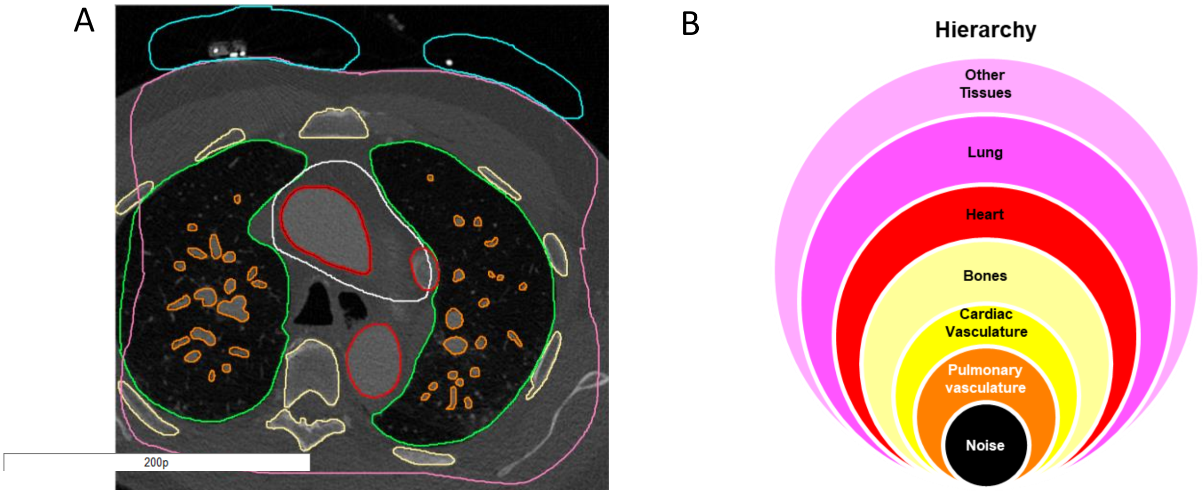


*Figure 26: Annotated chest CT scan. Nesting hierarchy for the large structures annotated: 'Non-tissue > Pulmonary vasculature > Major cardiac vessels > Bones > Heart > Lung > Other Tissues'.*
